# Orthogonal linear separation analysis: an approach to decompose the complex effects of a perturbagen

**DOI:** 10.1101/384446

**Authors:** Tadahaya Mizuno, Setsuo Kinoshita, Shotaro Maedera, Takuya Ito, Hiroyuki Kusuhara

**Affiliations:** Graduate School of Pharmaceutical Sciences, the University of Tokyo, Bunkyo-ku, Tokyo, 113-0033, Japan; ProMedico Co., Ltd., Ota-ku, Tokyo, 143-0023, Japan

**Keywords:** Connectivity Map (CMap), Factor Analysis, Mirror Data, OLSA, Profile Data

## Abstract

Drugs have multiple, not single, effects. Decomposition of drug effects into basic components helps us to understand the pharmacological properties of a drug and contributes to drug discovery. We have extended factor analysis and developed a novel profile data analysis method, orthogonal linear separation analysis (OLSA). OLSA contracted 11,911 genes to 118 factors from transcriptome data of MCF7 cells treated with 318 compounds in Connectivity Map. Ontology of the main genes constituting the factors detected significant enrichment of the ontology in 65 of 118 factors and similar results were obtained in two other data sets. One factor discriminated two Hsp90 inhibitors, geldanamycin and radicicol, while clustering analysis could not. Doxorubicin was estimated to inhibit Na^+^/K^+^ ATPase, one of the suggested mechanisms of doxorubicin-induced cardiotoxicity. Based on the factor including PI3K/AKT/mTORC1 inhibition activity, 5 compounds were predicted to be novel autophagy inducers, and other analysis including western blotting revealed that 4 of the 5 actually induced autophagy. These findings indicate the potential of OLSA to decompose the effects of a drug and identify its basic components. (<175 words)

## Introduction

The response to a drug can be a complex of the entire biological responses to the perturbagen and multiple responses in living systems. Not all the effects of a drug are fully discovered by researchers or developers. Therefore, to separate the complex effects of a drug into basic components is a prerequisite for a deep understanding of the pharmacological properties of drugs, which contributes to drug screening, drug repositioning, prediction of toxicity, and other properties.

Omics has made a great impact on biology since its emergence around the beginning of the 21^st^ century (Weinstein, 2001). Comprehensiveness of the methodology can translate the biological information of a sample into numeric data and, because of this characteristic, omics data are also called a profile. This quality of omics affords us mathematical approaches to comprehend the sample characteristics and are referred to as profile data analysis or, simply, profiling. To understand the complex effects of drugs, a substantial number of profiles have been accumulated and many methods of analysis have been devised, particularly for the mode of action (MOA) and toxicity (Andrusiak *et al*, 2012; Kim & Shin, 2014).

Of note, the Connectivity Map (CMap) project initiated by Broad Institute greatly contributed to the field (Lamb *et al*, 2006; Subramanian *et al*, 2017). In the project, dozens of microarray data analyzing cells treated with low molecular weight compounds were collected in the same platform. The concept is simple: a “signature” is simply defined by up- and downregulated genes responding to a perturbagen and the signatures can be compared to identify drugs with similar effects (Lamb *et al*, 2006). One of the essential features of this approach is not focusing on each gene but on the relationship of genes described as a gene pattern, or signature. There exist phenotypes that cannot be identified by the analysis of each gene (Mootha *et al*, 2003). Setting a novel layer with signatures, which effectively contract multiple variables of omics data and analyzing the signatures would enable the detection of such phenotypes that are difficult to recognize. Another curious characteristic of the CMap approach is that it does not depend on existing knowledge, which distinguishes this approach from gene ontology (GO) analysis or pathway analysis (Ashburner *et al*, 2000; Garcia-Campos *et al*, 2015). Utilization of existing knowledge in profiling is effective in reducing noise in profile data, while it restricts the capacity of analysis within the known. Analyses with CMap even utilize information unrecognized by researchers and therefore have the potential to show us the way to newly discovered findings. Many studies using CMap have succeeded in drug repositioning (Kosaka *et al*, 2013; Kunkel *et al*, 2011; Brum *et al*, 2015).

Considering the situation, we began to investigate whether it is possible to decompose the complex effect of a drug into basic components described by variable patterns using profile data analysis, particularly focusing on an unsupervised method. Among the matrix-decomposition methods, we focused on factor analysis (FA). FA decomposes a data matrix based on standard deviation, and is well established in various fields, including psychology and economics, and is also utilized in omics data analysis (Khan *et al*, 2014). Many studies accomplish dimension reduction and feature extraction of omics data to classify or investigate the similarity of samples with FA and other matrix-decomposition methods (Aziz *et al*, 2017; Zhang *et al*, 2012). However, to our knowledge, there are no studies that employ FA to separate the effects of a drug and extract the more basic components.

Among the several types of FA, the combination of principal component analysis (PCA) and following varimax rotation has been used extensively in the long history of FA. The characteristics are that the new indicators (factors in FA) composing the original variables are mutually orthogonal (Lever *et al*, 2017). We consider that the effect of a perturbagen can be described by a linear combination of more basic effects to some degree, while the remaining parts of the effect are nonlinearly integrated and not separable (Kundrát & Friedland, 2012). A strong point of the new indicators obtained by linear separation of an omics data matrix is that they are easier to comprehend than those obtained by nonlinear separation or machine learning (Gabel *et al*, 2014), which enables us to approach the molecular mechanism behind the composition.

A concern of utilizing FA with the principal component method in profiling is that the centroid in the novel coordinate space has no biological meaning and varies among data sets, which means that the obtained factors (vectors) in such a situation may not correspond to consistent biological meanings. To address that concern, we have extended FA by using “response profiles” and “mirror data” of the examined data set. When raw expression data of the treatment samples are normalized with the control data, the normalized data represent responses to the perturbagens and the centroid of that space is regarded as “no perturbation.” Assuming that the antagonizing or “reverse” responses exist, addition of the mirror data set to the original in PCA mathematically achieves an approximation of the novel coordinate space centroid to the data set in the original space. Therefore, when the procedure is applied to a response-profile data set, the generated space corresponds to the biological response space and the vectors constituting the space are considered to have consistent biological meanings. We call the profile data analysis with the simply modified FA, orthogonal linear separation analysis (OLSA). Here, we report the performance and possibility for OLSA to separate a perturbagen effect into basic components by analyzing transcriptome profiles and provide an easy-to-handle python scripts package for the novel method.

## Results

### The concept of orthogonal linear separation analysis of profile data

The workflow and concept of OLSA of a response-profile matrix are shown in Fig 1A, B, and C. Here, we define a “response-profile matrix” as a matrix with variables (e.g., gene expression change) in rows and samples (e.g., RNA-seq) in columns. An element in a response-profile matrix is a value representing a change of expression of a factor such as a log fold change or z score versus control. By converting the raw expression values of profile data into response values, the origin of the response data space represents the control treatment or no stimulation. One of the characteristics of OLSA is the use of a mirror data set (point-symmetric to the analyzed response-profile data). Considering the reversibility of biological responses, the mirror data set is a virtual data set that represents the antagonizing or reverse responses to the original data assumed to exist. FA of the combined data set mathematically enables approximation of the novel coordinate space centroid to the origin of the original data space where the variables are biologically relevant (here defined as the biological response space). Therefore, we can expect that the generated factors have consistent biological meanings. By employing OLSA, a response-profile matrix is described by the product of a response-vector matrix, a response-score matrix, and a total strength matrix, corresponding to the eigenvector matrix, the loading matrix, and a diagonal matrix of the L2-norm used for intensity correction (**Fig 1D**). FA with the principal component method describes an array of response-profile data with a linear combination of factors summarizing the original high-dimensional data, which helps us to comprehend the biological information of a profile data by investigating each separated factor (**Fig 1E**).

**Figure 1.**
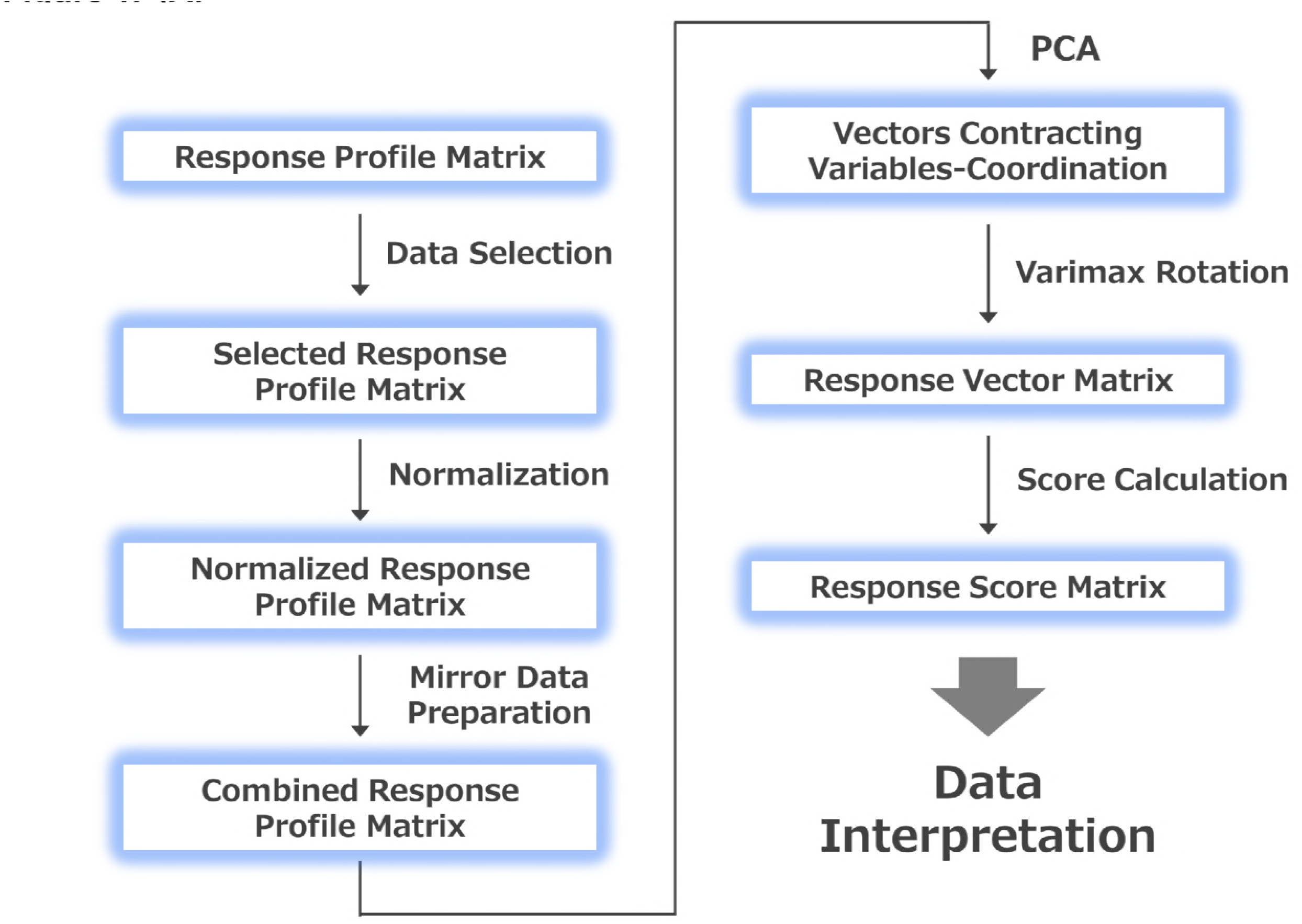
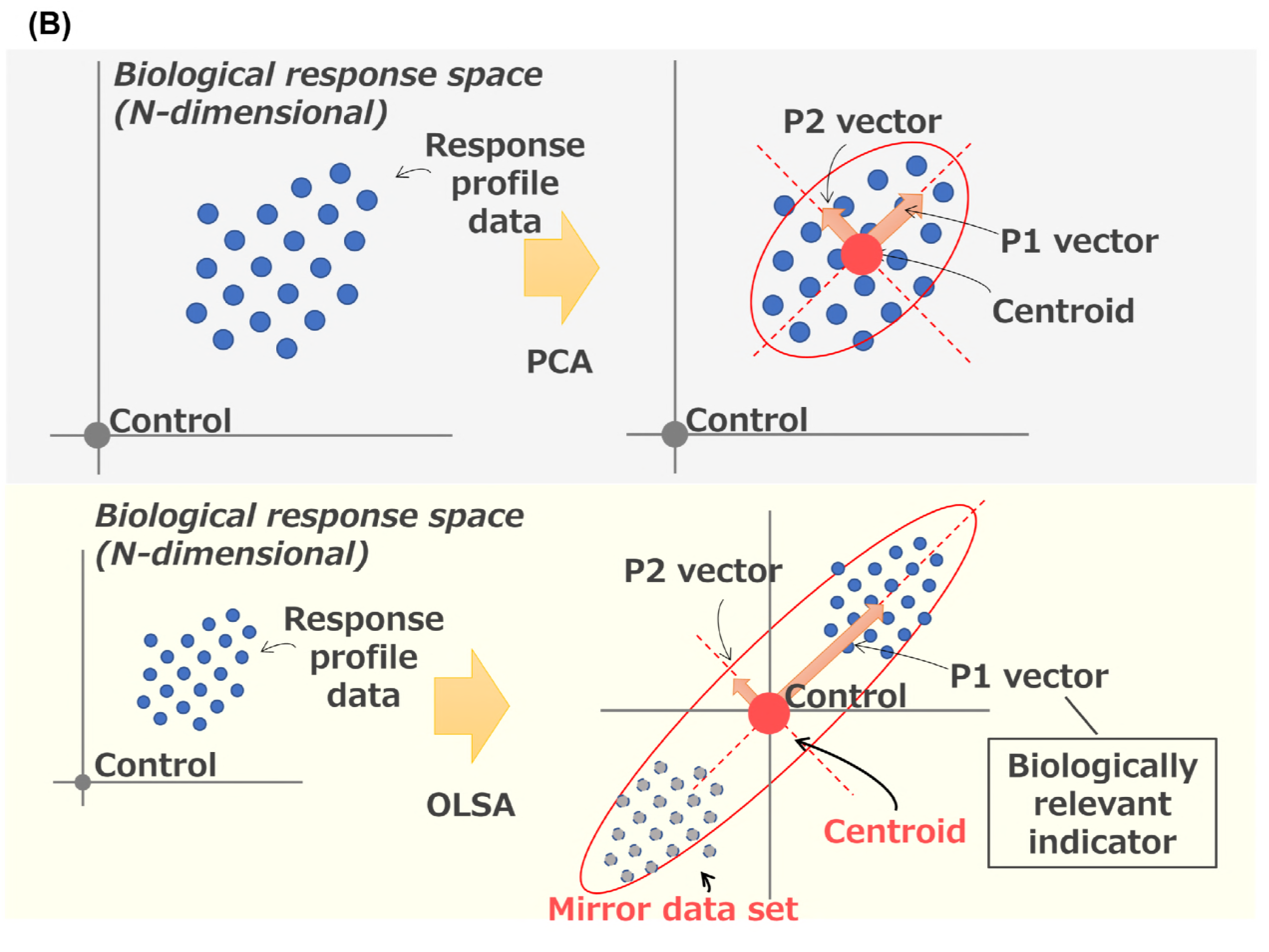
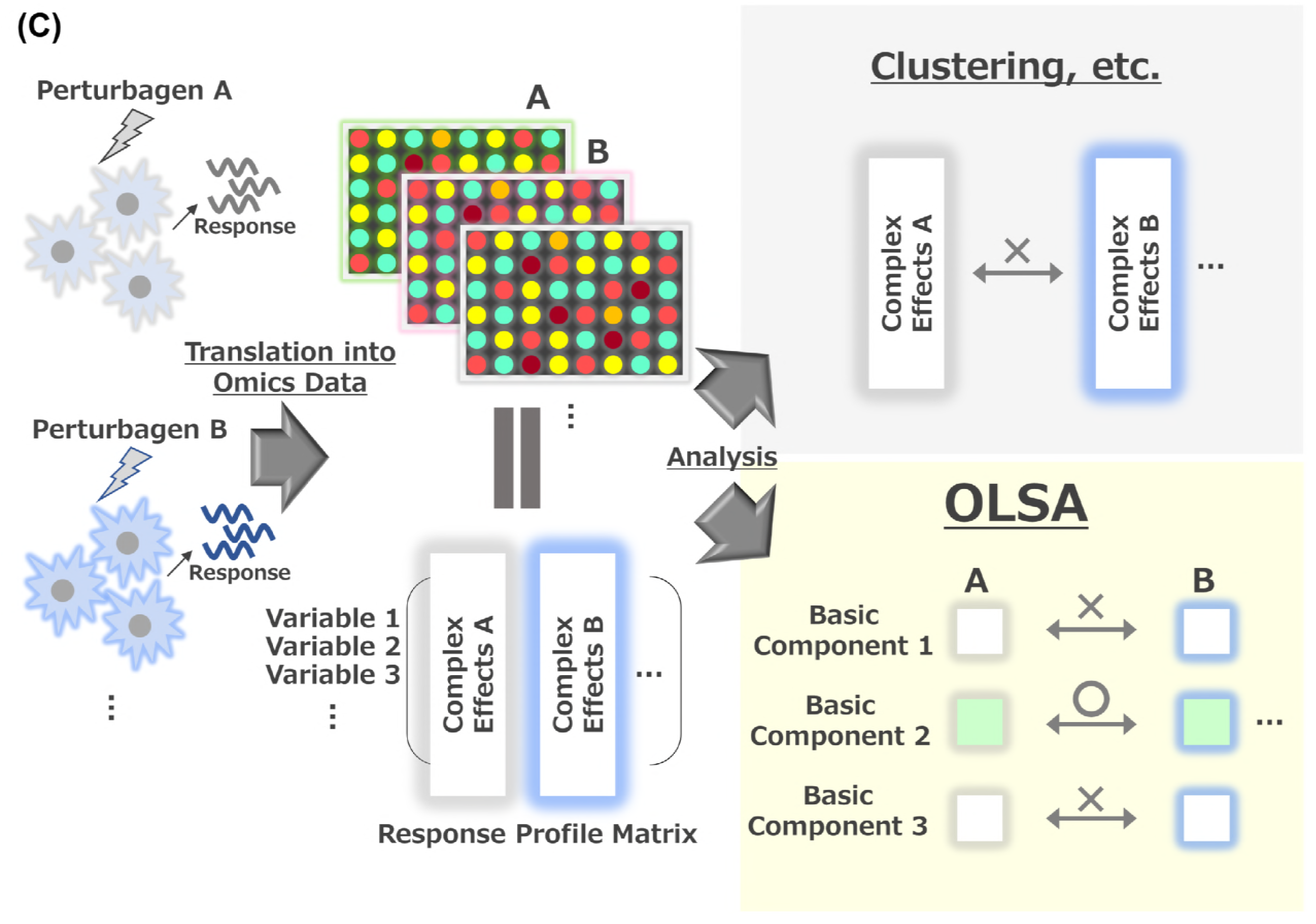
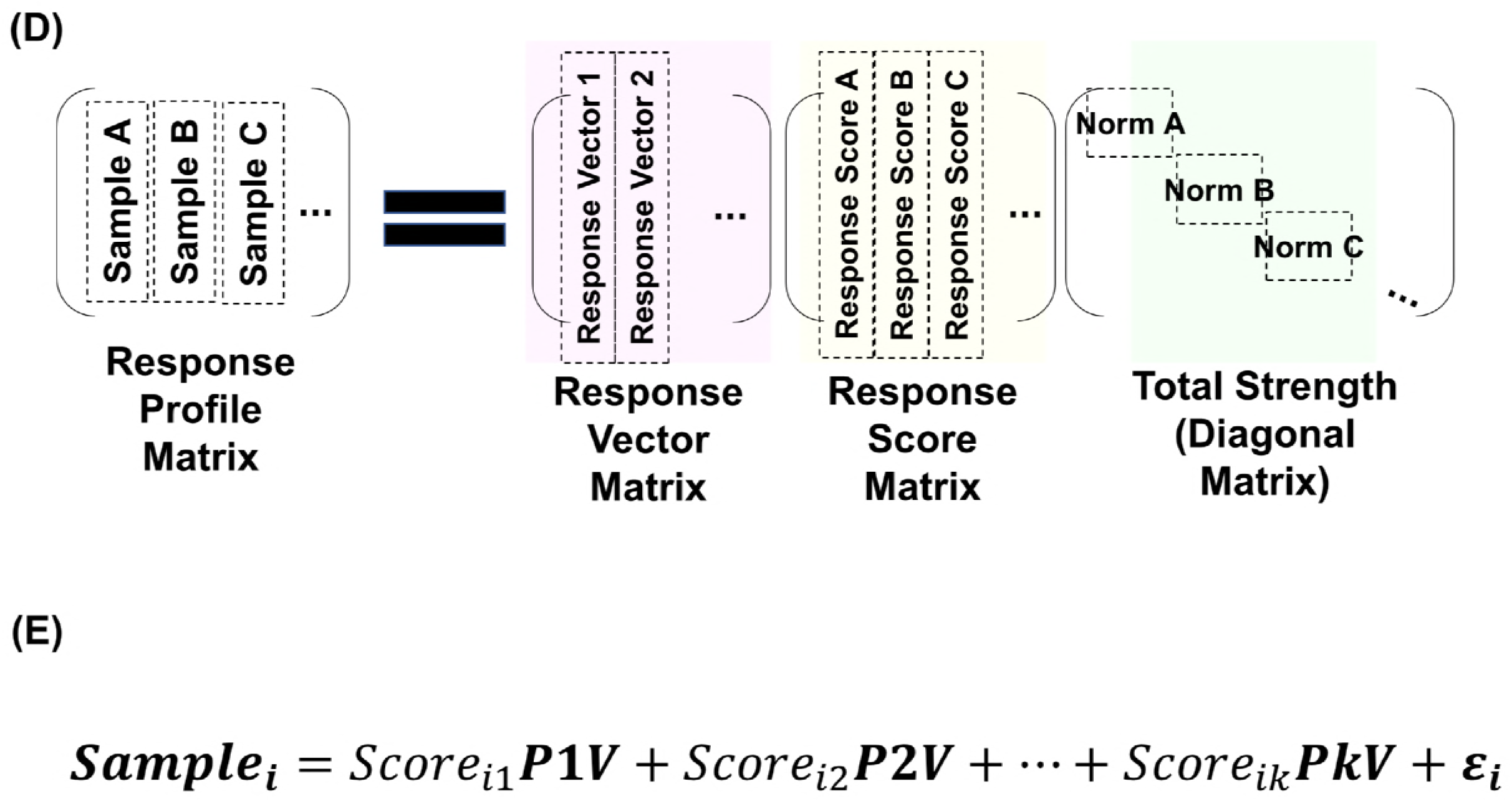
The concept of orthogonal linear separation analysis of profile data. A. Workflow of orthogonal linear separation analysis (OLSA). B. Illustration of OLSA concept. C. Illustration of OLSA application to response-profile data. D. Definition and relation of response-profile matrix, response-vector matrix, response-score matrix, and total strength. A response-score matrix is decomposed into the inner product of a response-vector matrix, a response-score matrix, and a diagonal matrix of the L2-norm. E. Description of a sample with a linear combination of the factors. When OLSA is applied to a response-profile data set with *p* variables in rows and *n* samples in columns, a sample is described in this way. *Sample_i_*, the data of ith sample where *i* = 1, &, n; *Score_ik_*, the response-score value of the kth response vector; *PkV*, the kth response vector; *ε_i_*, unobserved stochastic error terms.

We assumed that the factors isolated by OLSA are biologically relevant and the indicators of “basic” biological responses that constitute the original complex effect of a perturbagen. In the following, we confirmed that the factors generated with OLSA are biologically relevant and then investigated the application of the analysis method to understanding the effects of the perturbing drugs.

#### 1. Confirmation of biological meanings of the generated factors with OLSA Cellular responses in MCF7 cells treated with 370 perturbagens

We started with an analysis of the response-profile matrices obtained from CMap to verify OLSA. First, we analyzed the profile data set investigating the cellular responses of MCF7 cells treated with 318 compounds for 6 h as a training set (**Fig 2A, upper**). We subjected the data to varimax rotation and analyzed 118 vectors, accounting for 80% of cumulative contribution (**Fig 2B**). To gain insight into the biological relevance of the factors obtained, we used the GO analysis of large weight genes in a focusing factor. Genes constituting a factor were sorted by their contribution to the factor and the top 1 % of genes were subjected to GO analysis. There were 65 factors with significant enrichment of GO and the ratio of such factors (hereafter termed significant enrichment of GO ratio, SEGR) to the total was 0.551 (65/118) (**Fig 2C**).

**Figure 2.**
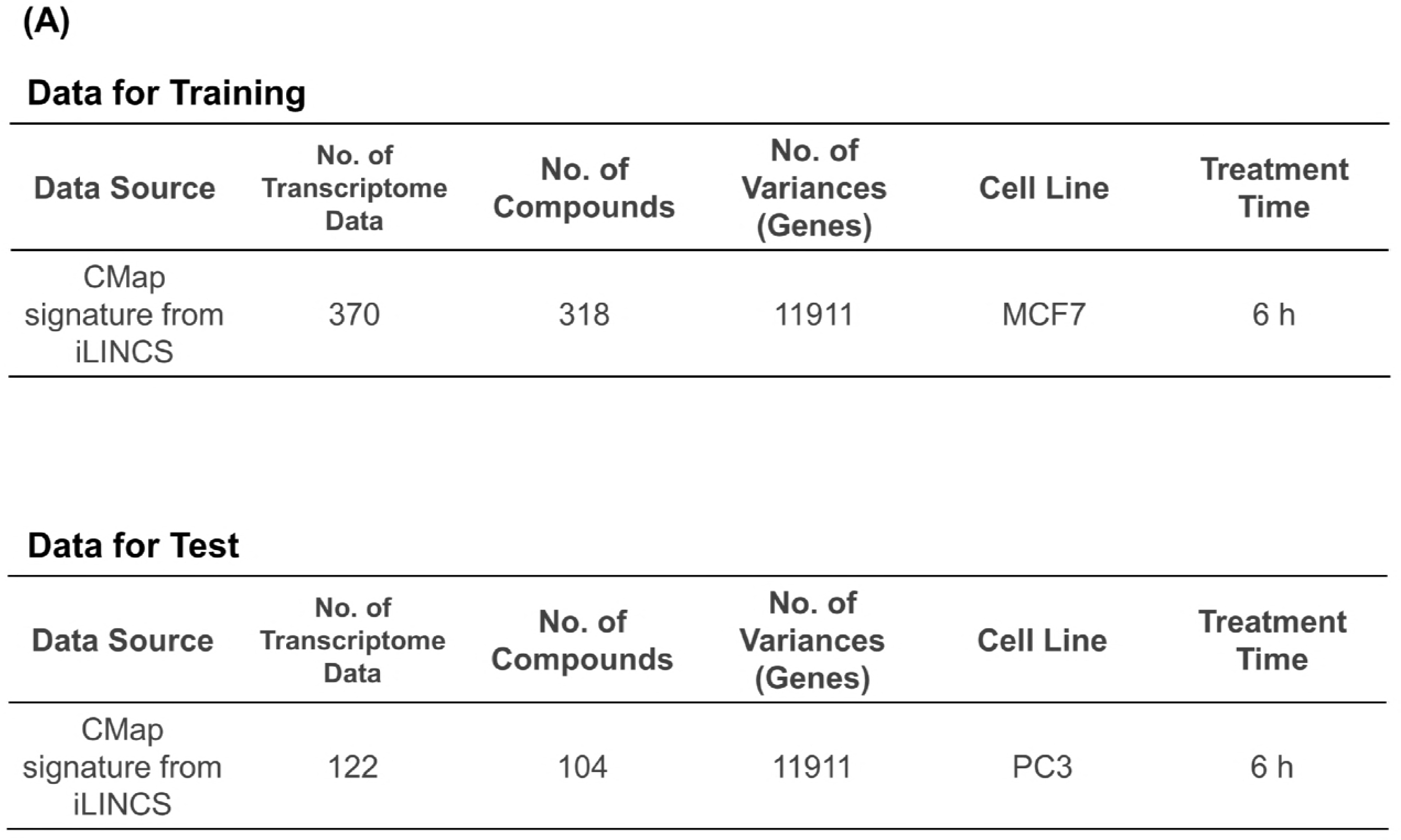
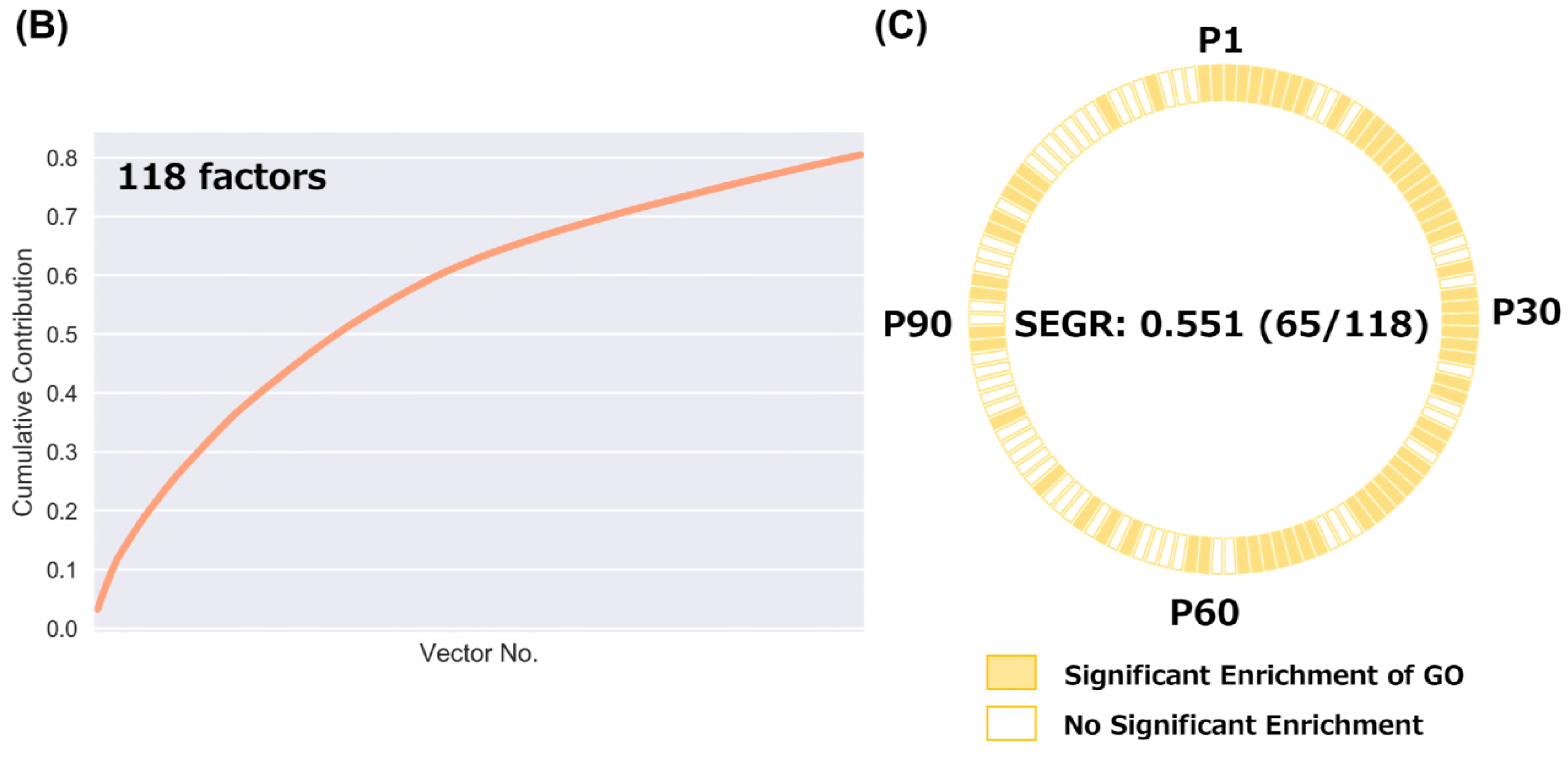
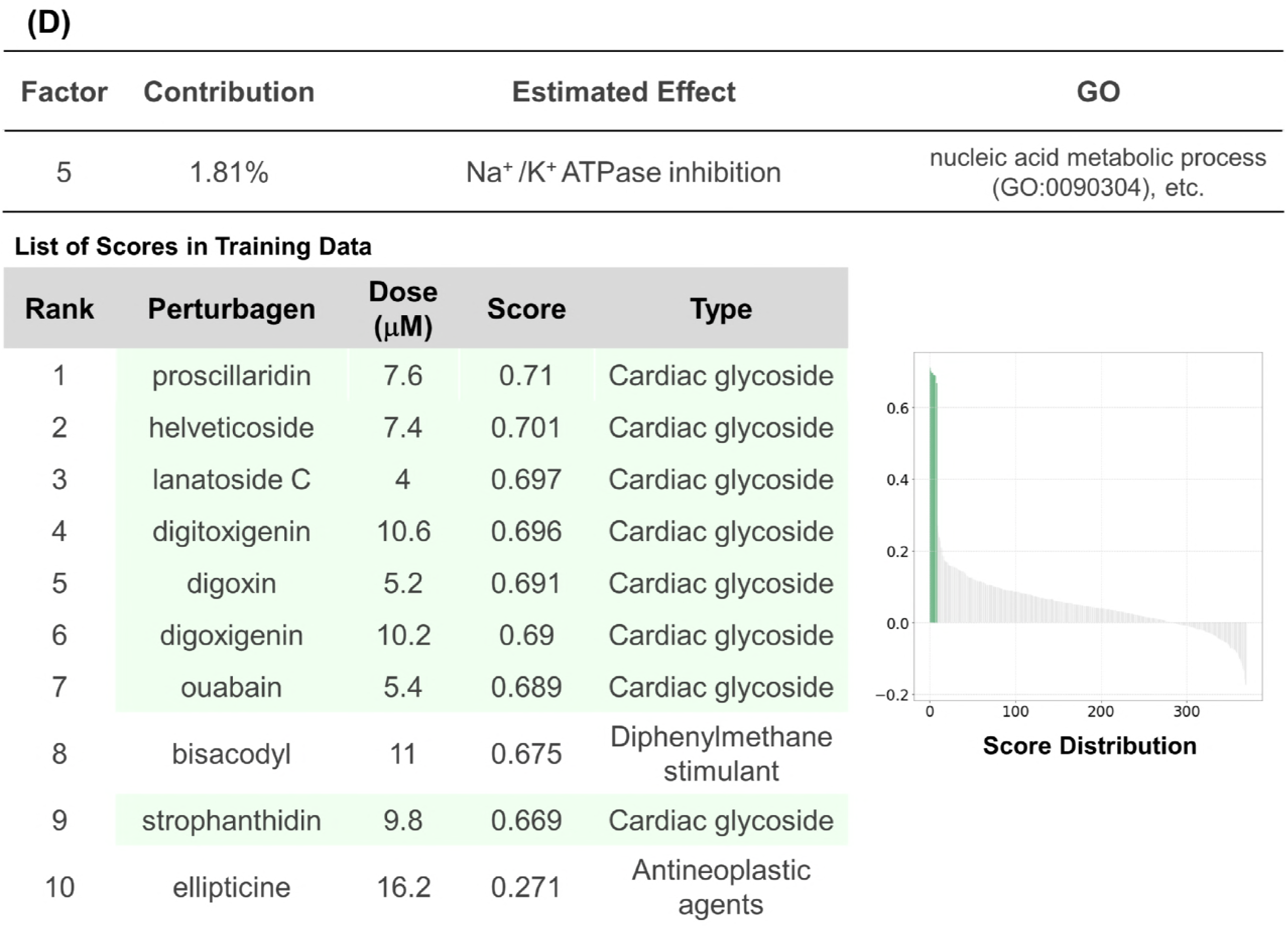
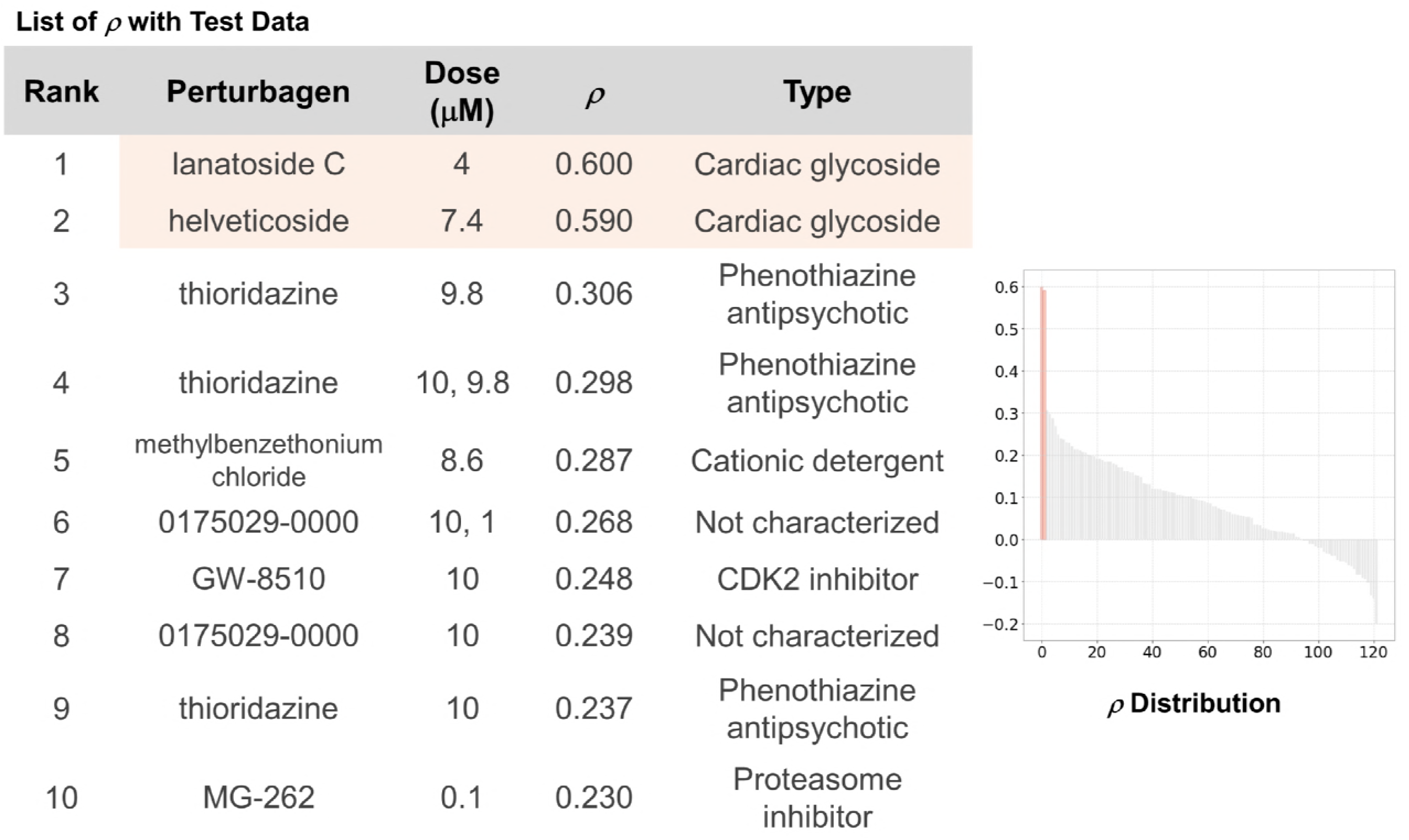
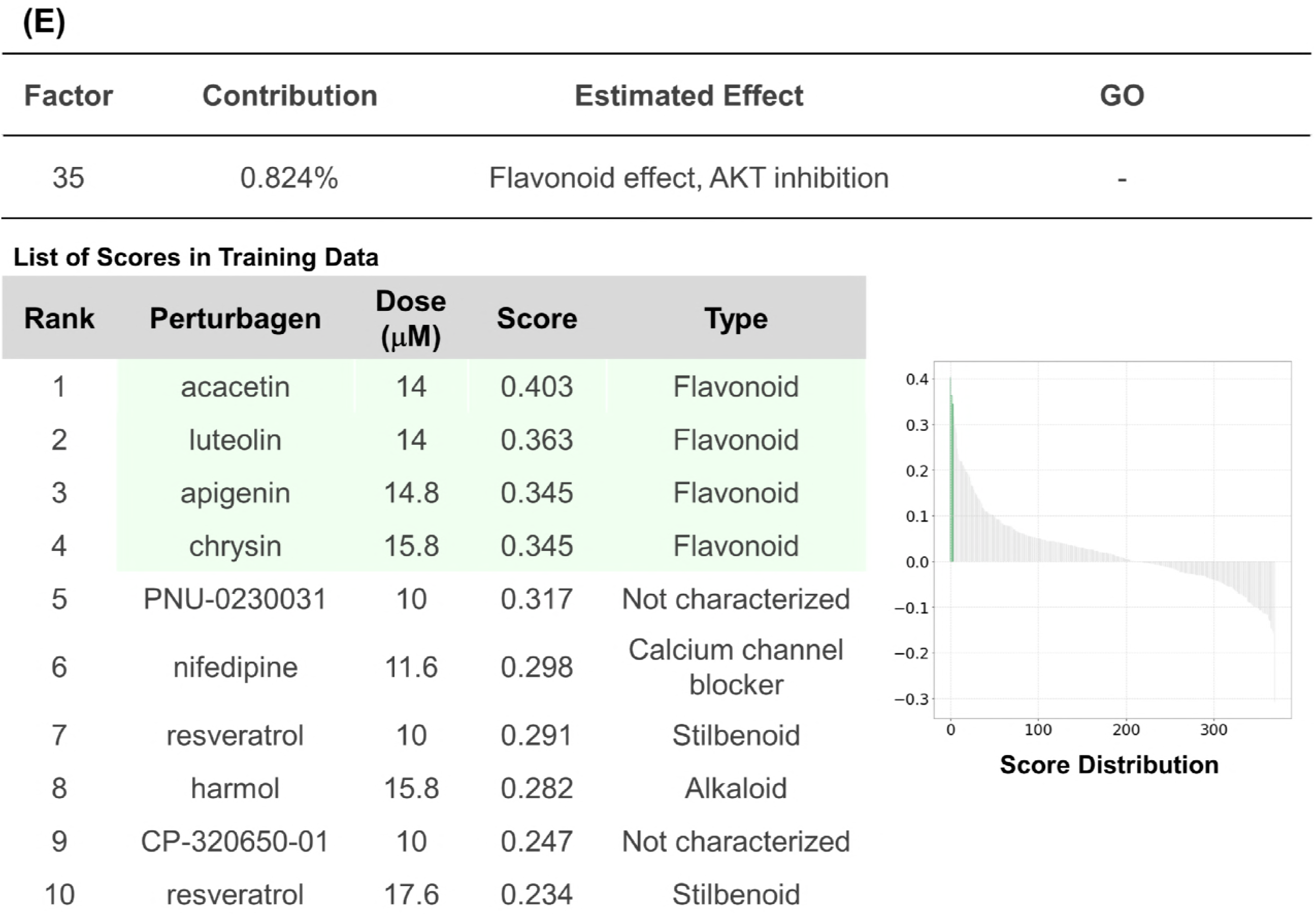
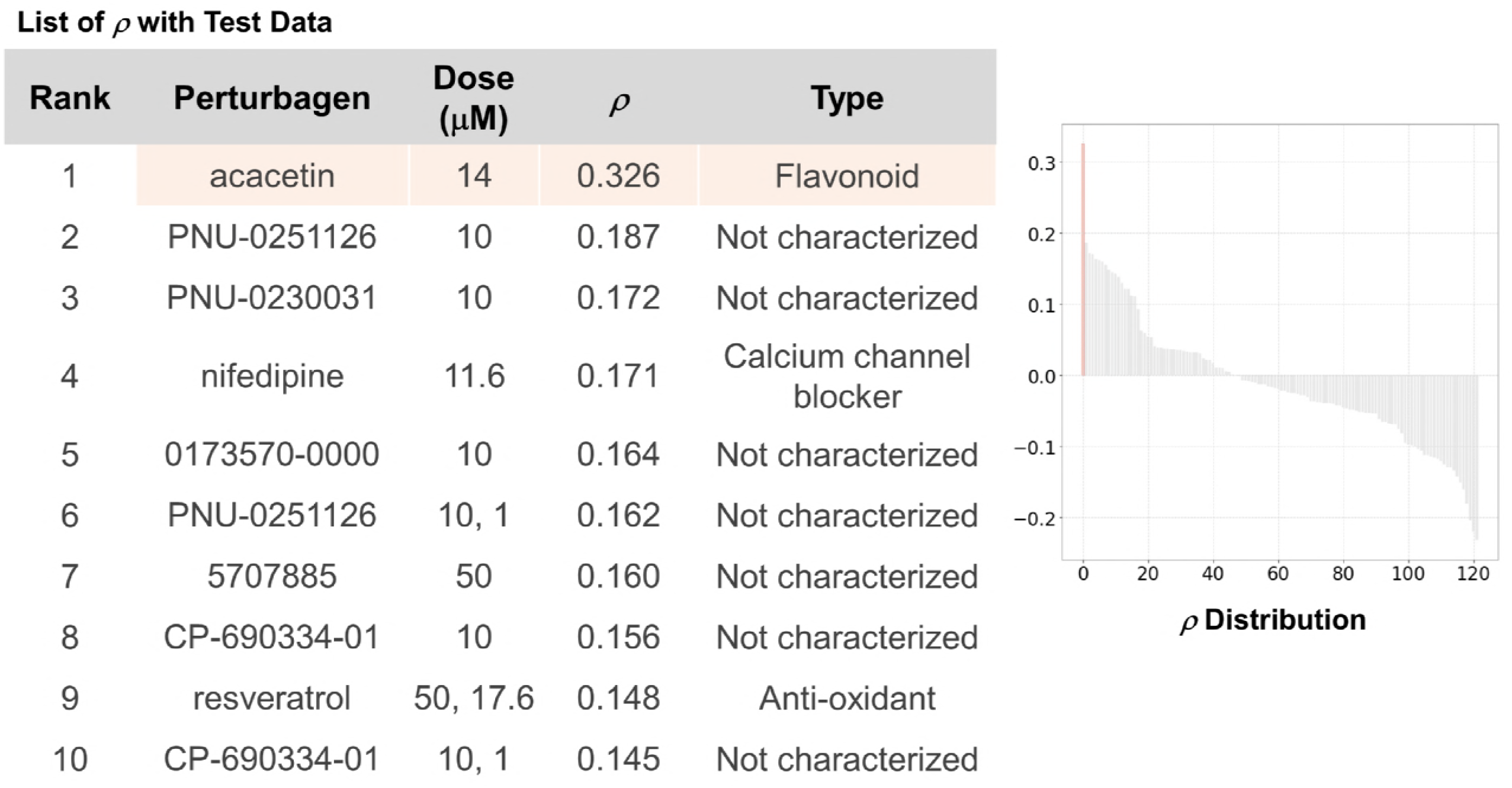
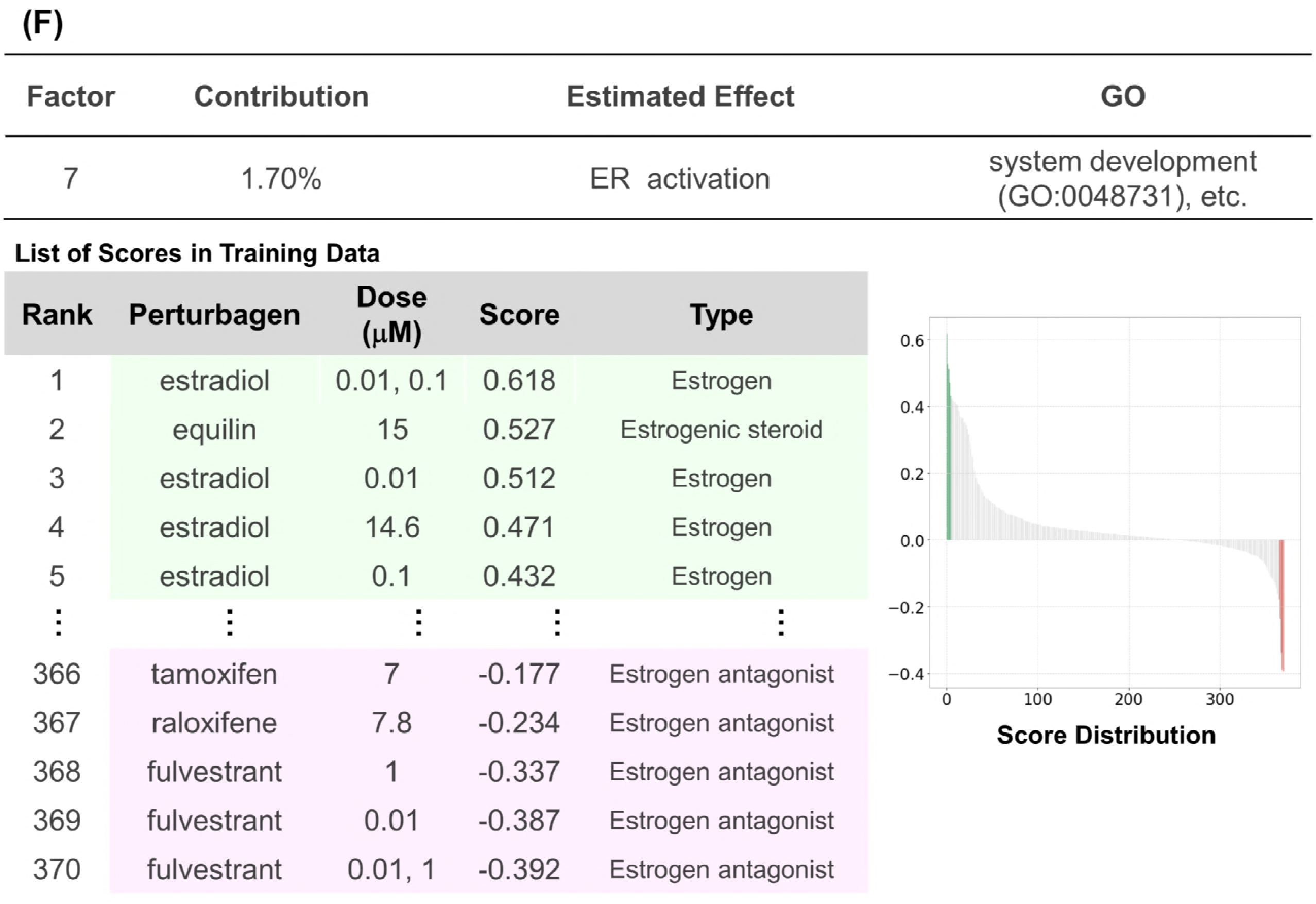
Analysis of cellular responses in MCF7 cells treated with 370 perturbagens. A. Data sets employed in this figure. B. The cumulative contribution curve of the factors contracting the training data set. The contribution of each factor to the total deviation was calculated and arranged in descending order. The cumulative contribution was calculated from the top and plotted. C. Plot of the factors whose main constituents exhibit significant enrichment of gene ontology. Genes constituting a response vector were sorted by the square of each value. The top 1% of genes were subjected to GO (biological process) analysis using Enrichment analysis of the Gene Ontology Consortium. Factors annotated with significant enrichment of GO after multiple-testing corrections (Benjamini-Hochberg method, α < 0.05) was depicted in yellow-filled squares. SEGR, significant enrichment of GO. D. Analysis of P5 factor. P5 factor (the factor with the 5^th^ highest contribution) scores and rho (*p*) of all compounds are arranged in descending order and plotted on the “Score Distribution” graph and *“ρ* Distribution” in each data set, respectively (upper, training; lower, test). Green or light salmon in the graph indicates “cardiac glycoside.” The rank, name, dose, and score of the top 10 compounds are shown. E. Analysis of the P35 factor using the method described in D. F. Analysis of the P7 factor. Green and red in the graph indicate estrogens and antiestrogens, respectively. The rank, name, dose, and score of the top and last 5 compounds are shown.

Given that the generated factors are biologically relevant, the gene patterns are precise signatures, supposed to be conserved in another data set, and useful in identifying compounds with a focused cellular response. To verify this supposition, we characterized several factors using a detailed literature survey and subsequently calculated Spearman’s correlation between the selected factors and another data set. For the test data set, we employed a profile set comprising 122 transcriptome data analyzing PC3 cells treated with 104 compounds provided by CMap (**Fig 2A, lower panel**).

The high-scoring compounds in the factor with the 5^th^ highest contribution (hereafter P5 factors) were cardioglycosides and all 8 cardioglycosides in the data set were ranked in the top 9 (**Fig 2D, upper panel**). Notably, the 8^th^ compound, Bisacodyl, is used as a stimulant laxative drug, but is reported to inhibit Na+/K+ ATPase (Schreiner, 1980), consistent with the cardioglycoside target. In addition, both cardioglycosides in the test set, lanatoside C and helveticoside, were ranked in the top 2 of the compound list sorted by Spearman’s correlation coefficients, supporting that the P5 factor includes cardioglycoside effects, such as Na+/K+ ATPase inhibition (**Fig 2D, lower panel**).

As for P35 factor, although the gene pattern exhibited no enriched GO, flavonoids with a similar structure were enriched in the top four: acacetin, luteolin, apigenin, and chrysin (**Fig 2E, upper panel and Fig EV1A**). One of the cellular responses they have is AKT inhibition (Kim *et al*, 2014; Lim *et al*, 2016; Kim & Lee, 2018; Yang *et al*, 2014). It is consistent that the other compounds around the top of the list, nifedipine (6^th^), resveratrol (7^th^ and 10^th^), and harmol (8^th^), are potent inhibitors of AKT (Kaimoto *et al*, 2010; Haider *et al*, 2005; Abe & Kokuba, 2013). Acacetin, the only flavonoid with a focused structure in the test set, exhibited the highest correlation coefficient, and nifedipine and resveratrol were also ranked at the 4^th^ and 9^th^ positions in the list of 122 compounds, respectively (**Fig 2E, lower panel**).

Interestingly, the literature survey suggested that the P76 factor was associated with ion modulation responses although the factor contribution was quite low (0.3% of the total, Fig EV1B). P7 factor also had a curious characteristic: the positive high-scoring compounds in the factor were estrogens, while the negatively scoring compounds were antiestrogens (**Fig 2F**). Although we were not able to validate the characteristic in another data set because of lack of estrogen data in the test set, it was consistent with the result from CMap analysis and suggests that the signs of the response scores correspond to the direction of basic cellular responses (Lamb *et al*, 2006). Together, these data support that factors separated linearly by OLSA reflect cellular responses in the CMap data set.

### Cellular responses in HepG2 cells treated with genotoxic compounds

To investigate whether OLSA works in data sets other than CMap, we applied the method to data obtained from the public transcriptome database, GEO (https://www.ncbi.nlm.nih.gov/gds). Magkoufopoulou *et al* investigated the transcriptome profiles of HepG2 cells treated with 158 genotoxic compounds and obtained 474 transcriptome data (Magkoufopoulou *et al*, 2012). We employed these data and separated them into two groups: the data of 24 h treatment for training and the data of 12 and 48 h for the test (**Fig 3A**). The data in each set were aligned and converted into two response-profile matrices by eliminating missing values and calculating robust *z* scores for the control treatment data, and the processed training data set was subjected to OLSA. The analysis generated 29 factors from 186 transcriptome data up to 80% cumulative accumulation (**Fig 3B**). It should be noted that the cellular responses of virtually 62 perturbagens are summarized into 29 factors because each perturbagen had 3 biological replicates, which may explain the apparently small number of factors compared with the number of data. GO analysis revealed that 21 of 29 (SEGR; 0.724) factors exhibited significant enrichment of GO (**Fig 3C**).

**Figure 3.**
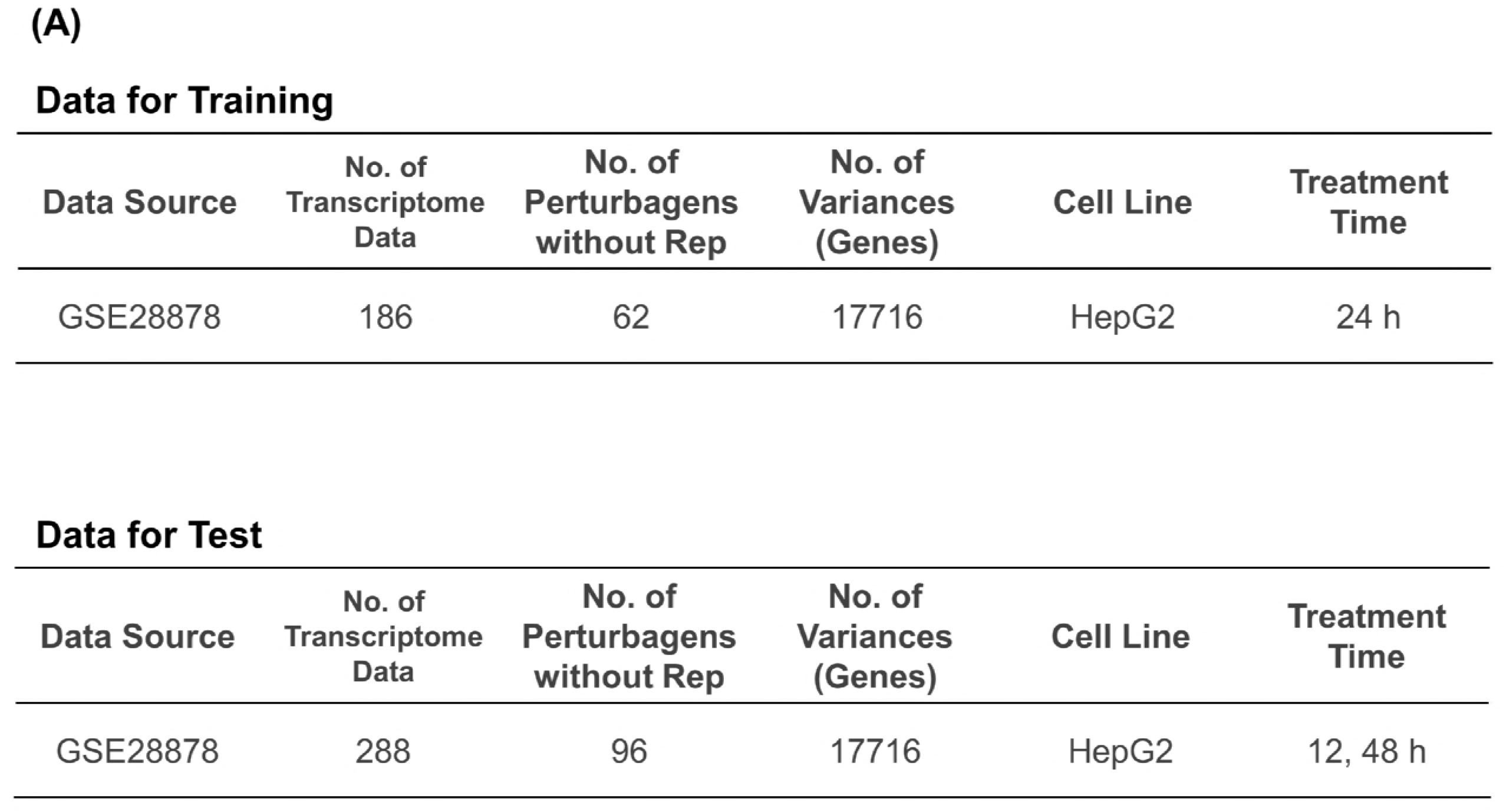
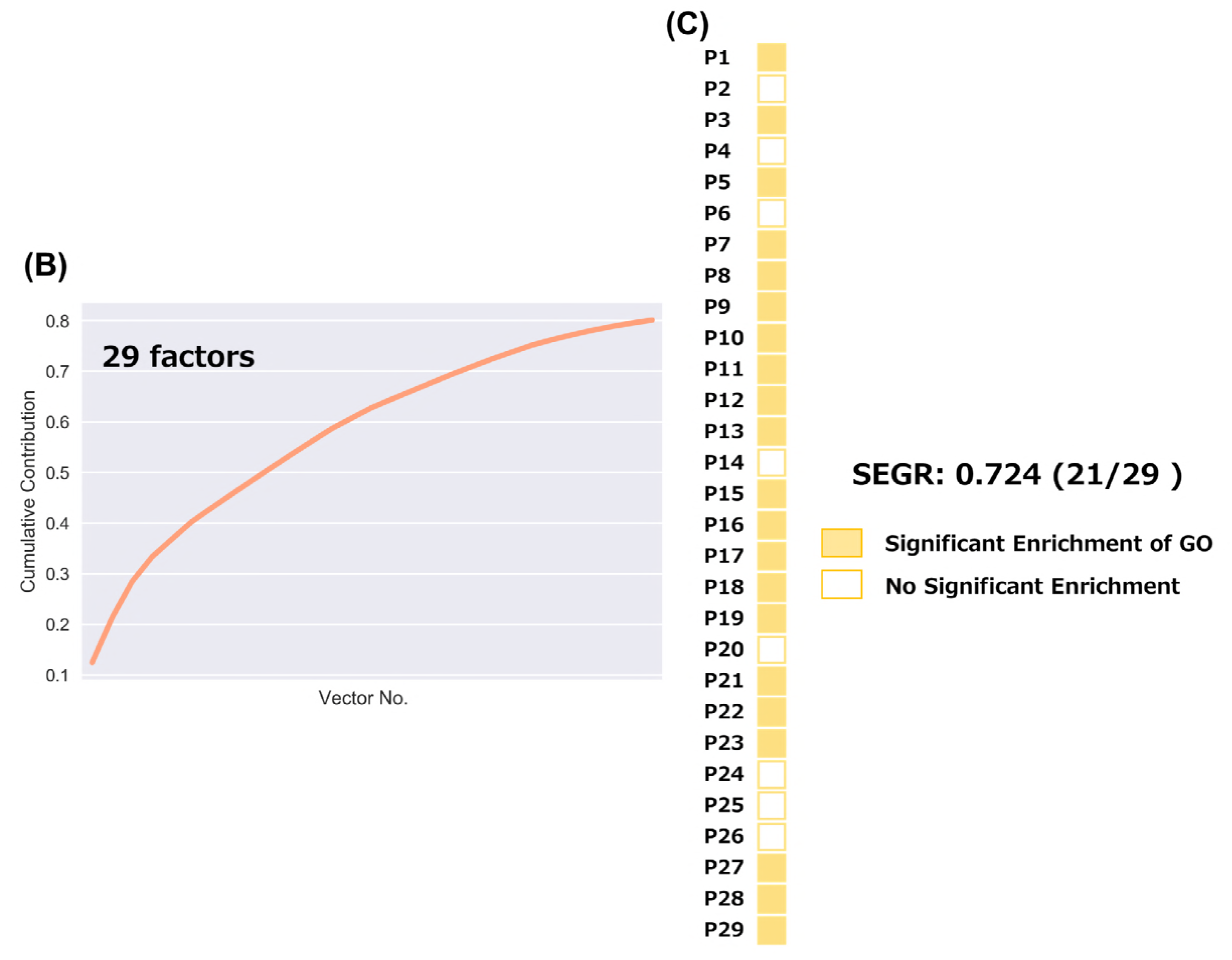
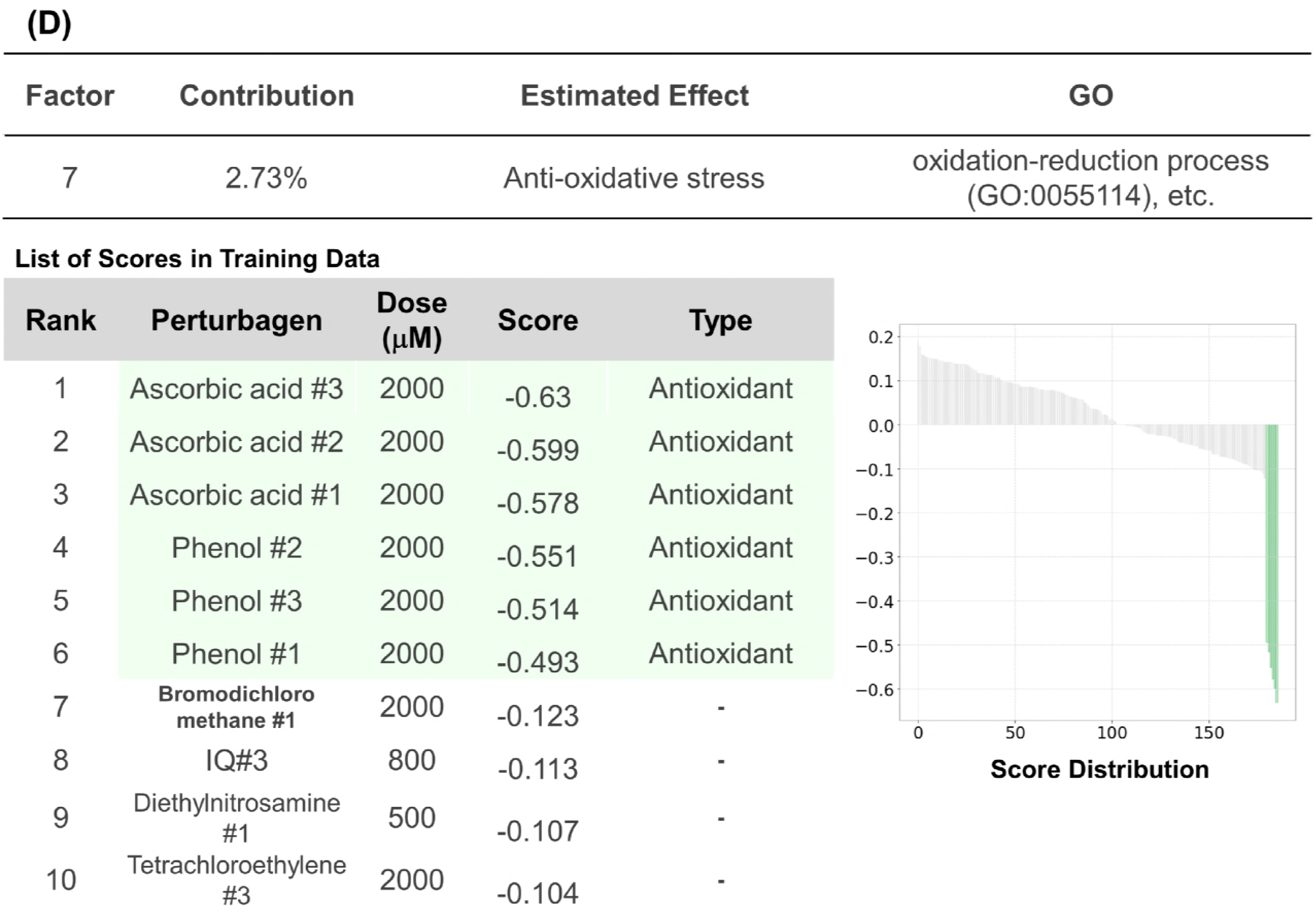
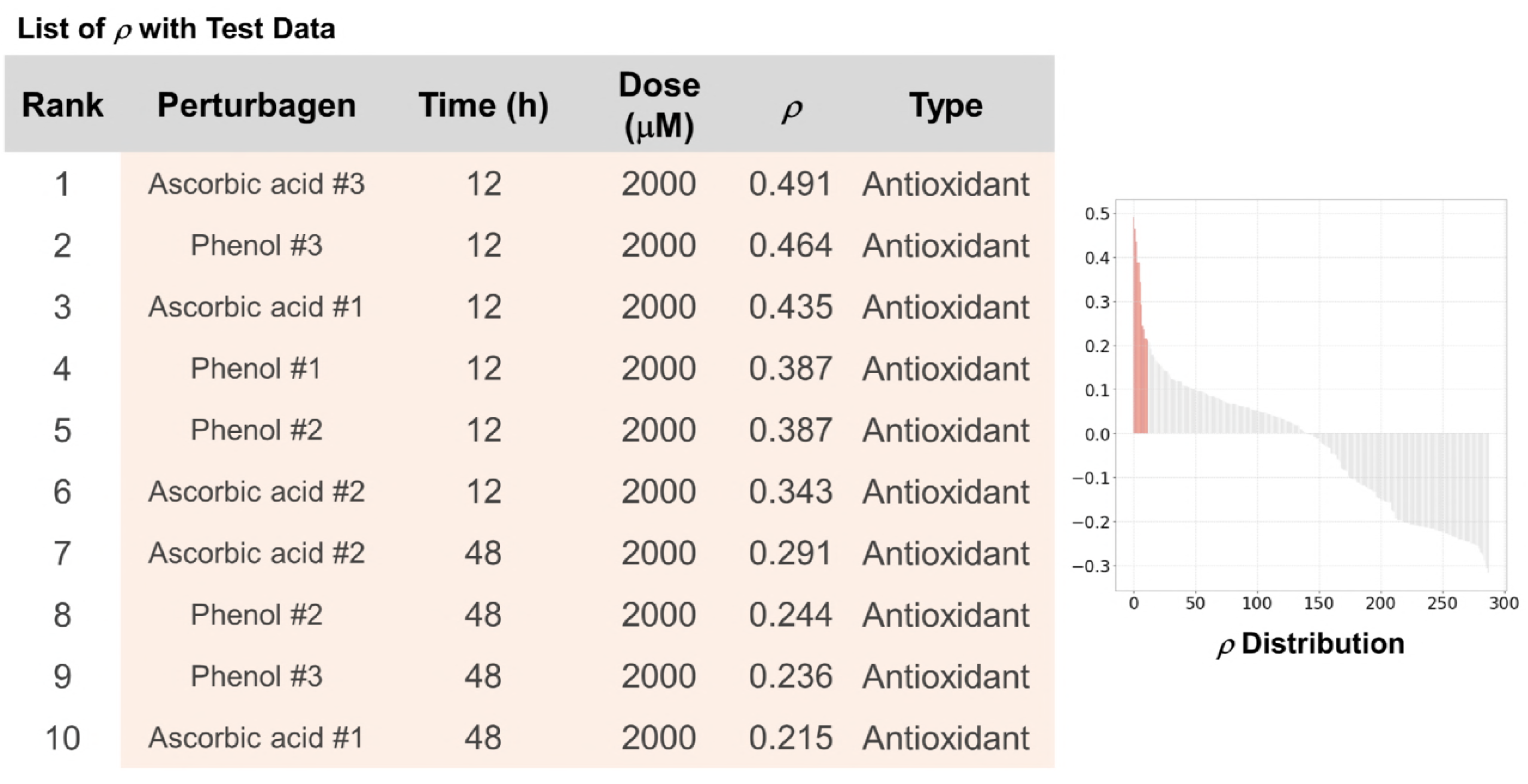
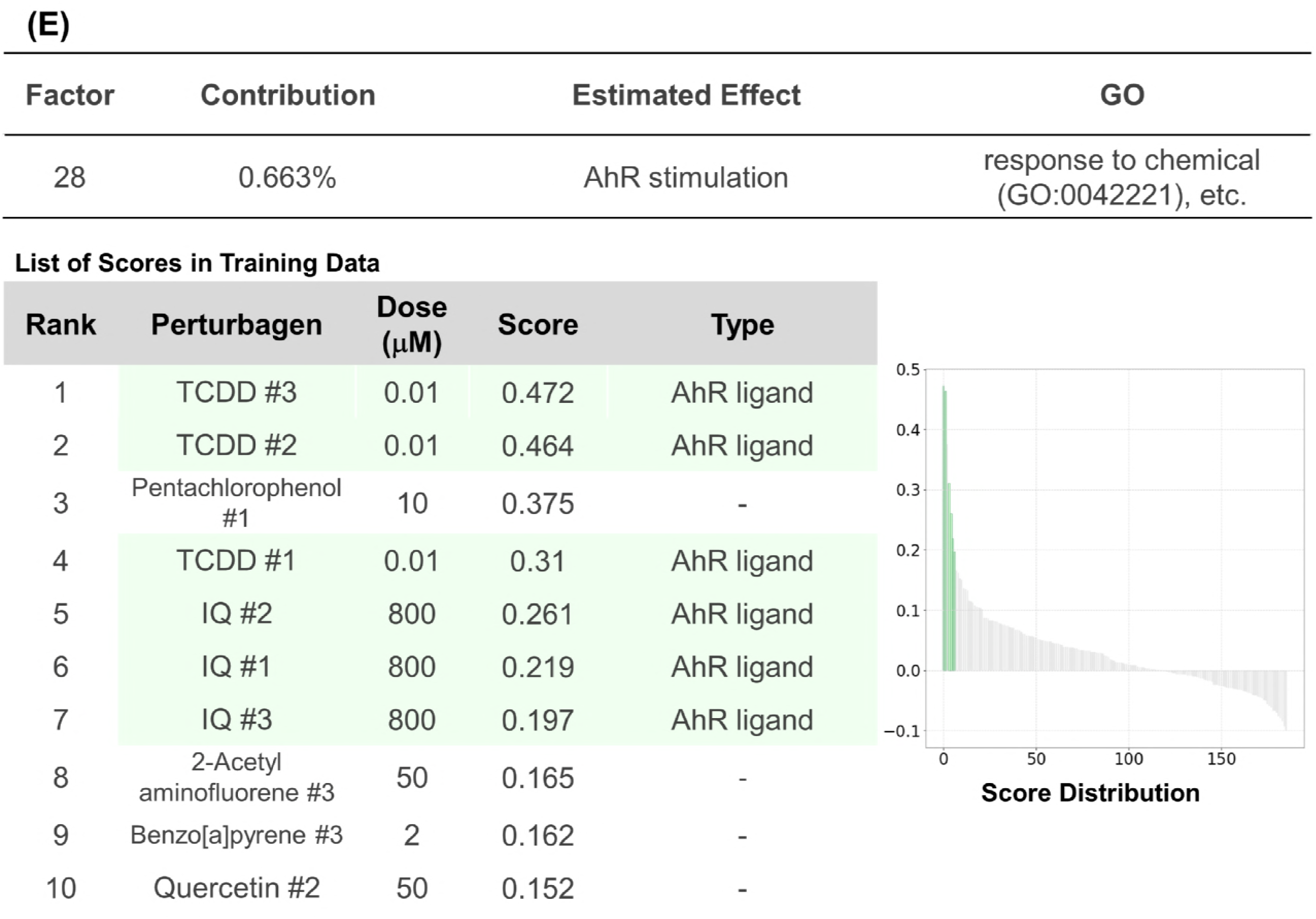
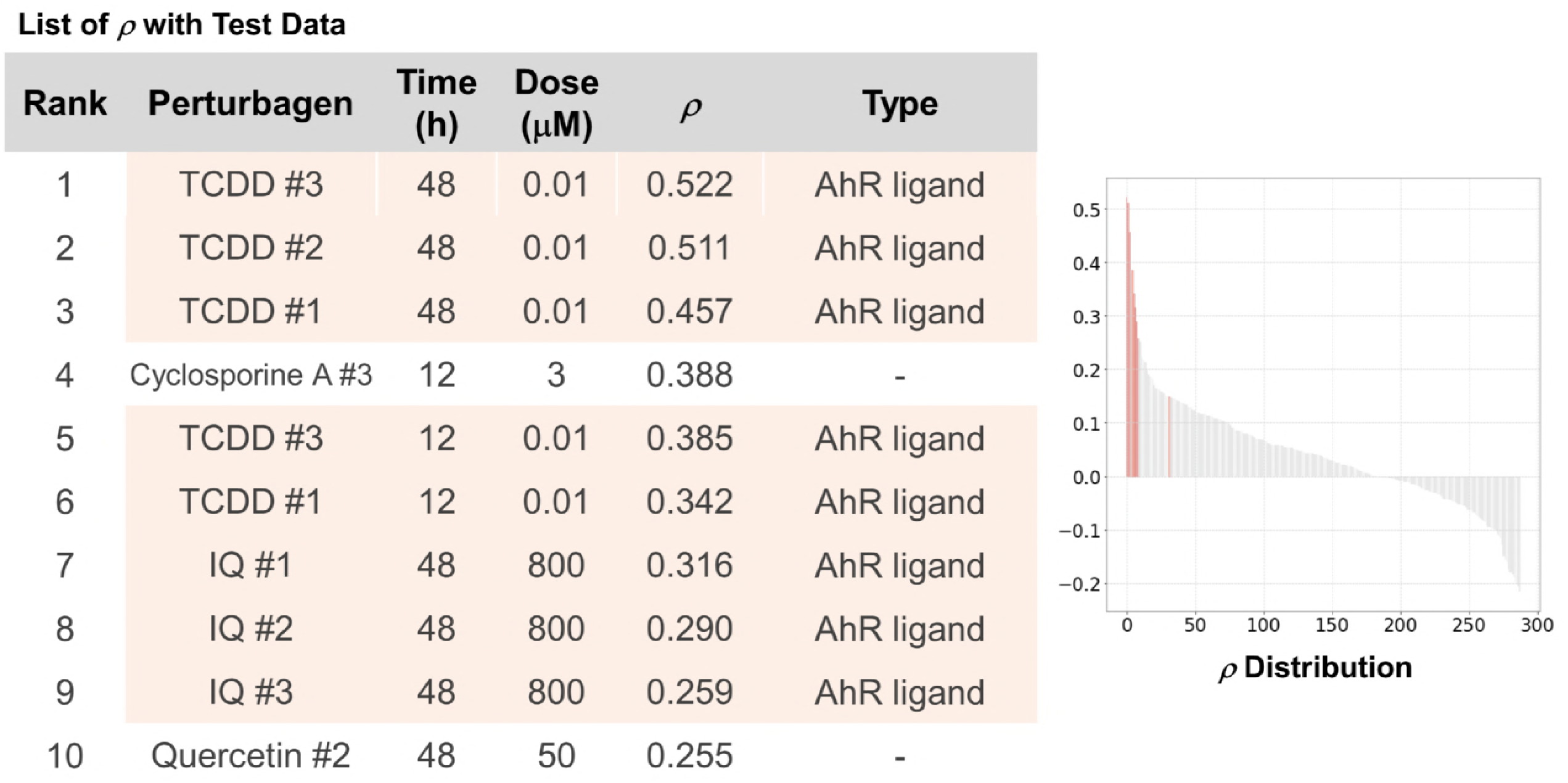
Analysis of cellular responses in HepG2 cells treated with 62 genotoxic compounds. A. Data sets employed in this figure. B. The cumulative contribution curve of the factors comprising the training data set. The contribution of each factor to the total deviation was calculated and arranged in descending order. The cumulative contribution was calculated from the top and plotted. C. Plot of the factors whose main constituents exhibit significant enrichment of gene ontology. Genes constituting a response vector were sorted by the square of each value. The top 1% of genes were subjected to GO (biological process) analysis using Enrichment analysis of Gene Ontology Consortium. Factors annotated with significant enrichment of GO after multiple-testing corrections (Benjamini-Hochberg method, α < 0.05) are depicted in yellow-filled squares. SEGR, significant enrichment of GO. D. Analysis of P7 factor. P7 factor scores and rho (*p*) of all compounds are arranged in descending order and plotted on the “Score Distribution” graph and *“ρ* Distribution” in each data set, respectively (upper, training; lower, test). Green or light salmon in the graph indicates ascorbic acid or phenol, respectively. The rank, name, dose, and score of the top 10 compounds are shown. “-” and “#” indicate not investigated in the literature survey and the number of biological replicates, respectively. E. Analysis of the P28 factor using the method described in D. Green or light salmon in the graph indicates TCDD or IQ, respectively. TCDD, 2,3,7,8-tetrachlorodibenzo-p-dioxin; IQ, 2-amino-3-methylimidazo[4,5-*f*]quinolone.

In a detailed investigation of individual factors, several factors were clearly connected to biologically relevant responses. For instance, the negative-scoring compounds in the P7 factor were dominated by ascorbic acid, one of the GOs significantly enriched in the factor is “oxidation-reduction process (GO:0055114),” and phenol and both of them had antioxidant properties in common (**Fig 3D, upper panel**) (Foti, 2007). The test set was validated by calculating Spearman’s correlation coefficients between the gene pattern and the test set. Vitamin C and phenol exhibited high values regardless of treatment time, supporting the consistency of the biological meaning of the factors.

Aryl hydrocarbon receptor (AhR) is a transcription factor belonging to the bHLH-PAS family and coordinates the transcripts for xenobiotic responses (Stockinger *et al*, 2014).2,3,7,8-Tetrachlorodibenzo-*p*-dioxin (TCDD), a representative dioxin and AhR ligand, was ranked in the top 4 of the compound list sorted by P28 factor scores (**Fig 3E, upper panel**). In the same list, the top 5^th^, 6^th^, and 7^th^ ranked perturbagen compounds were biological replicates of 2-amino-3-methylimidazo[4,5-*f*]quinolone (IQ) and IQ is reported to be an AhR ligand also, which implies that the P15 factor includes an AhR response (Sekimoto *et al*, 2016). Consistent with this, significant enrichment of “response to chemical (G0:0042221)” was observed by GO analysis of the P28 factor constituents. High Spearman’s correlation coefficients were calculated between the gene pattern and the test data set derived from TCDD- or IQ-treated cells and these support that the P28 factor has consistent biological roles (**Fig 3E, lower panel**).

In the same way, the P15 and P29 factors were suggested to include P450 modulation and interferon I stimulation effects, respectively (**Fig EV2A and B**). These results indicate that OLSA application is not restricted to well-aligned data sets such as those provided by CMap.

### Inflammatory responses in macrophages

We investigated the capacity of OLSA in a response-profile matrix composed of relatively few data. Inflammatory responses are strictly regulated by many factors, for example, the types of stimulants and time courses (Di Gioia & Zanoni, 2015). Raza *et al* investigated transcriptional networks in murine macrophages treated with several inflammatory stimulants at various time points by analyzing the transcriptome data set composed of 60 data with 30 perturbagens (2 biological replicates each) and we employed this data set (Raza *et al*, 2014). First, the data set was separated into training and test sets: the data of bone marrow derived macrophages (BMDM) from BALB/c mice for the training and BMDM from C57/BL6 mice for the test (**Fig 4A**). It should be noted that the training set contained 3 different stimulants, lipopolysaccharide (LPS), interferon β, and interferon γ, while the test set comprised only LPS-treated BMDM data. The training data set was processed to obtain a response-profile matrix in a manner similar to that with Magkoufopoulou’s data and subjected to OLSA. We obtained 15 factors and the SEGR was 0.33 (5/15) (**Fig 4B and C**).

**Figure 4.**
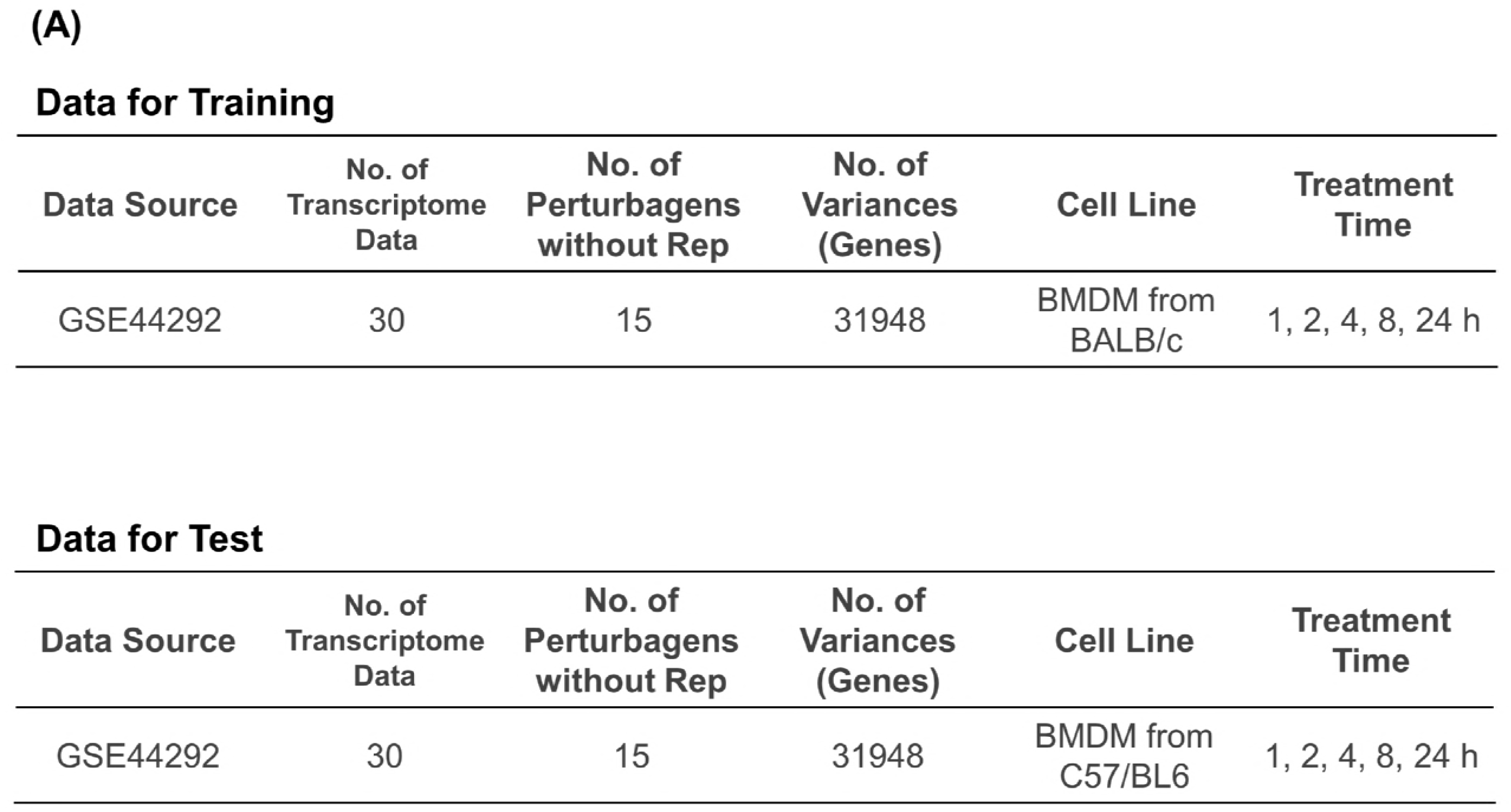
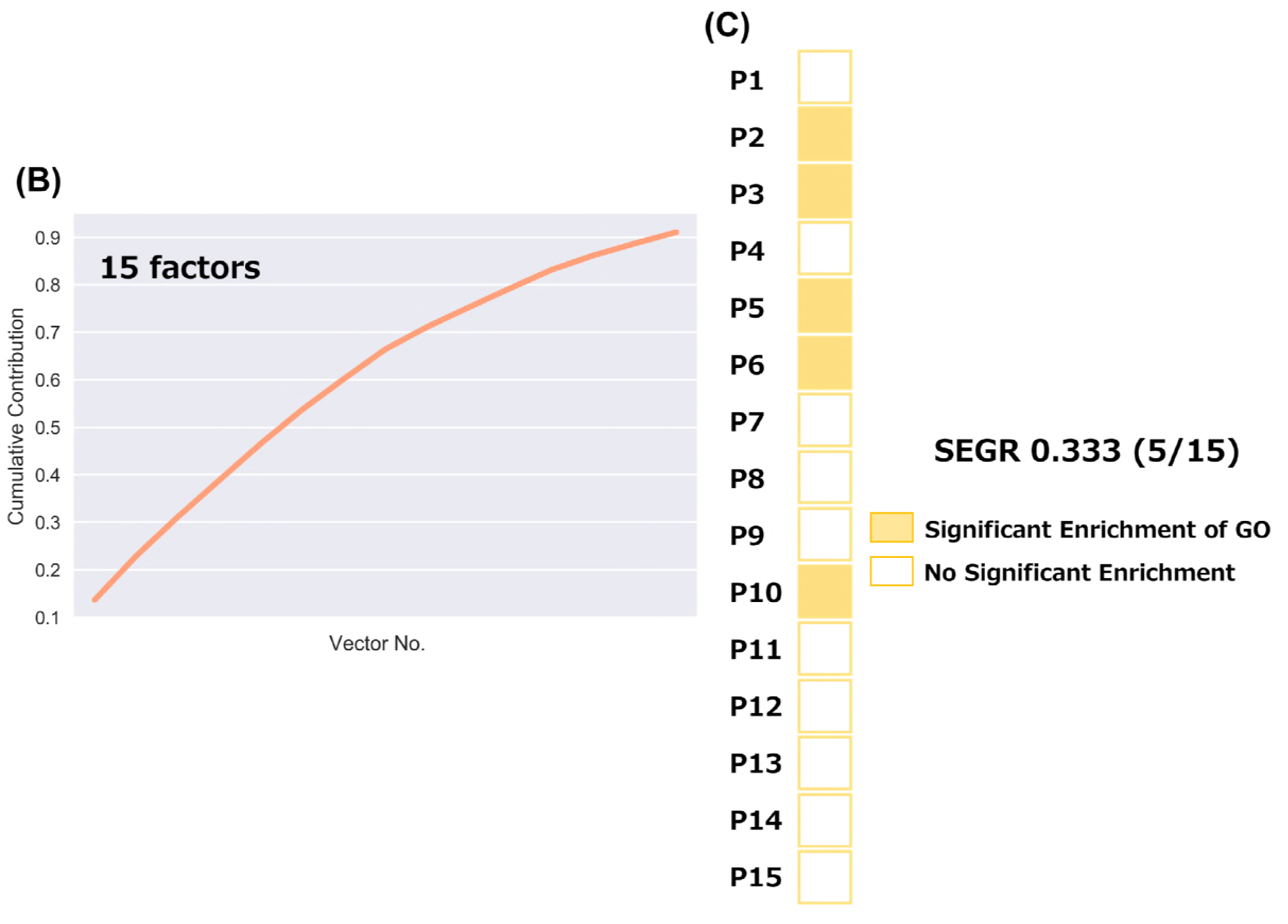
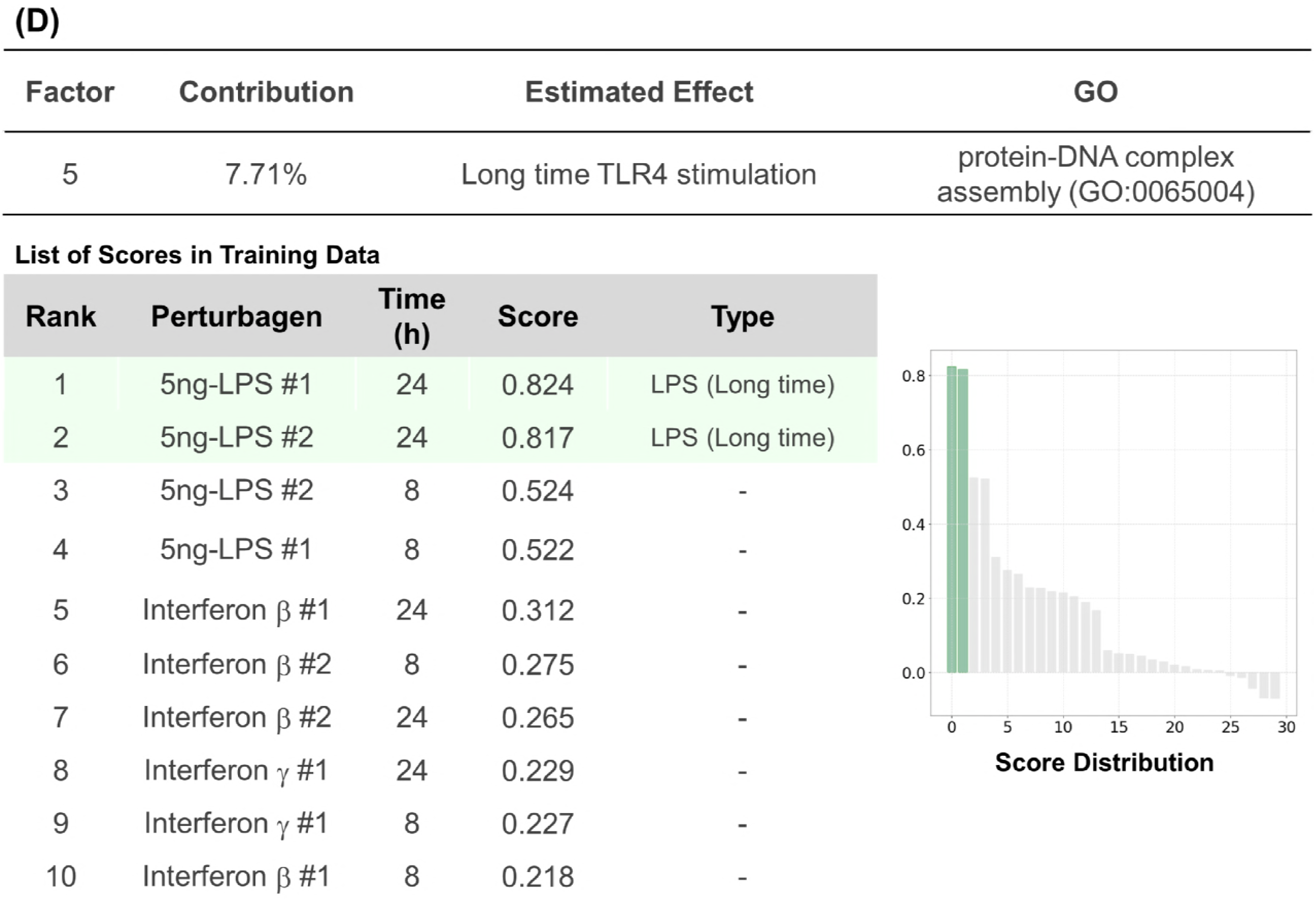
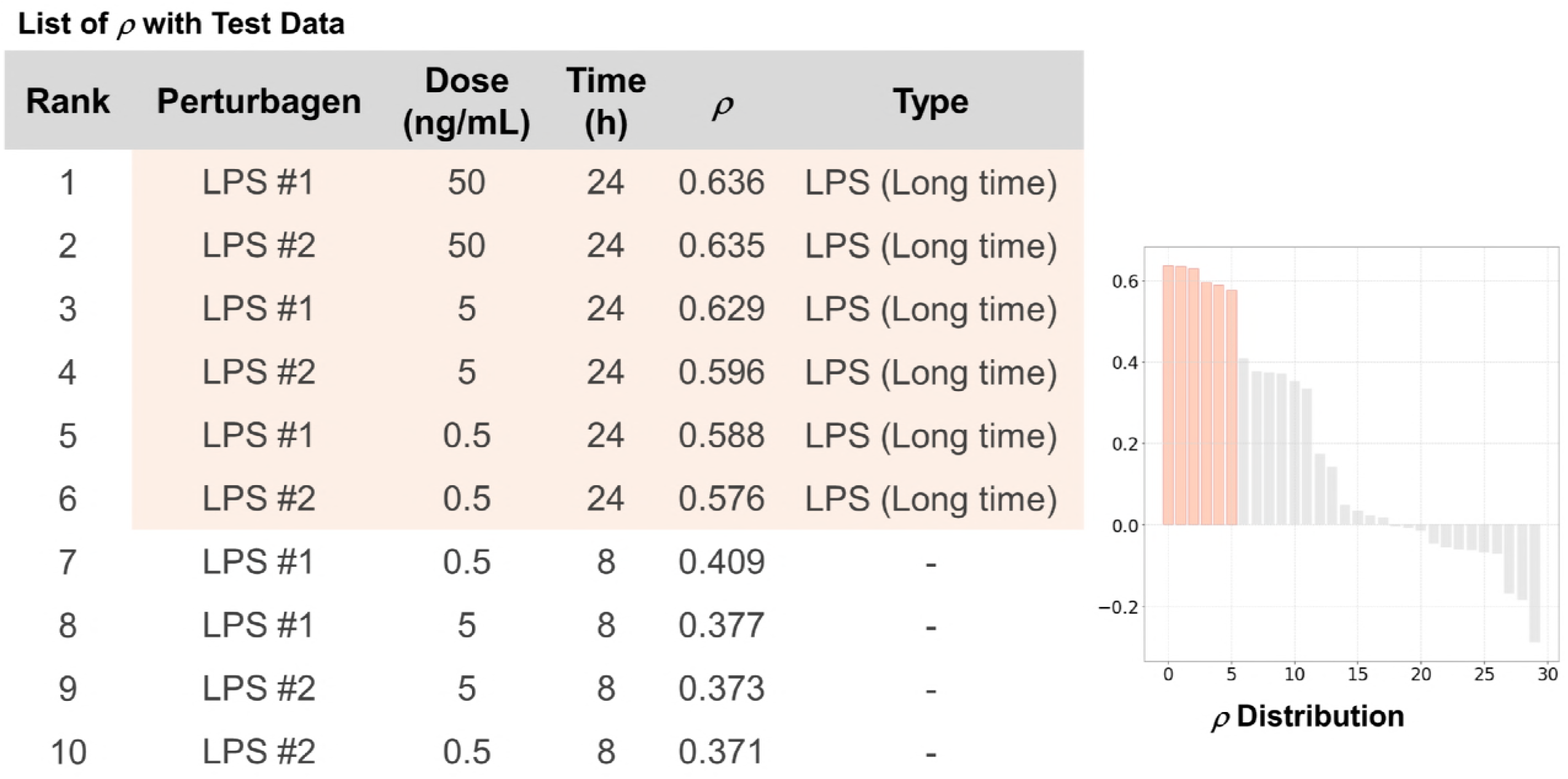
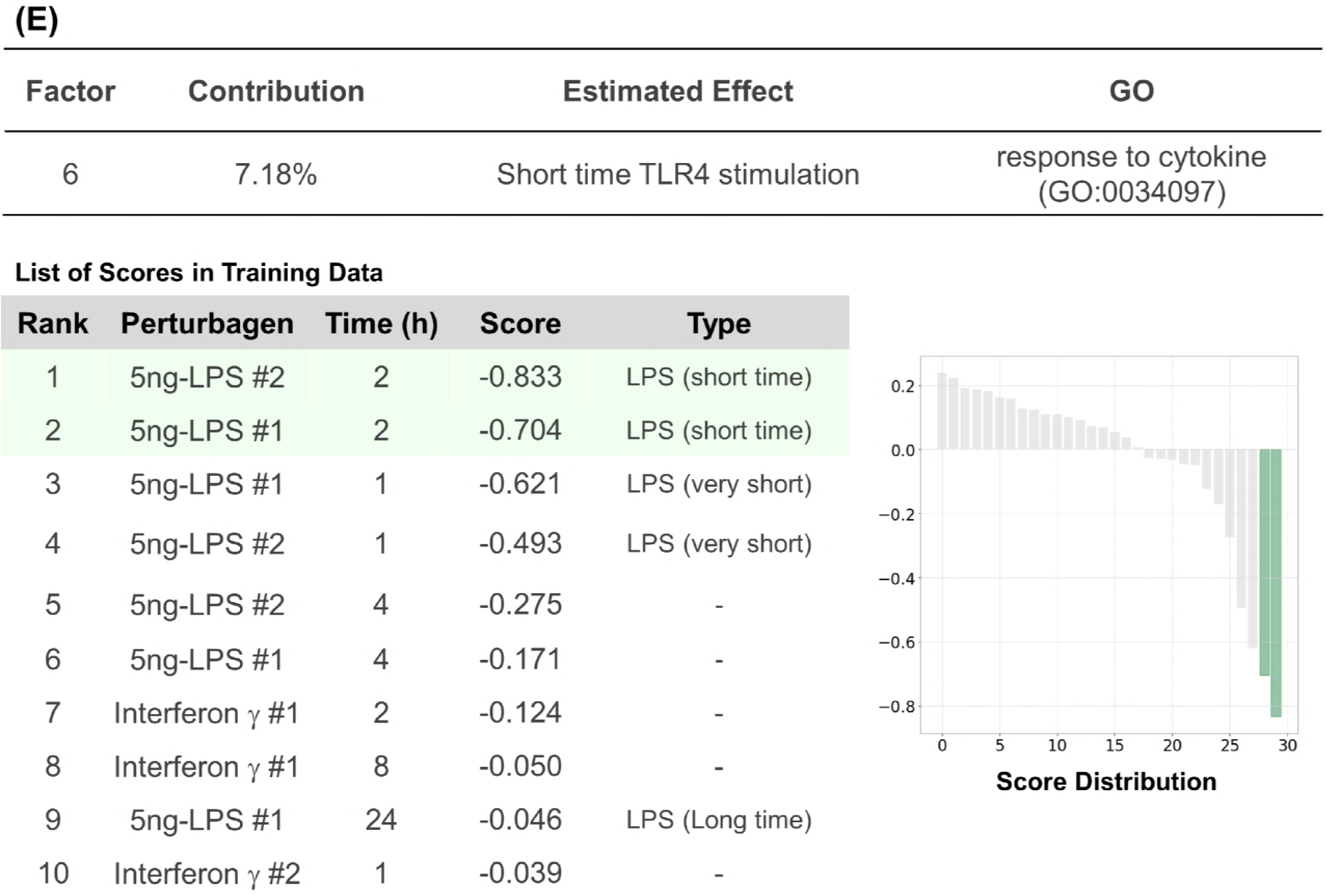
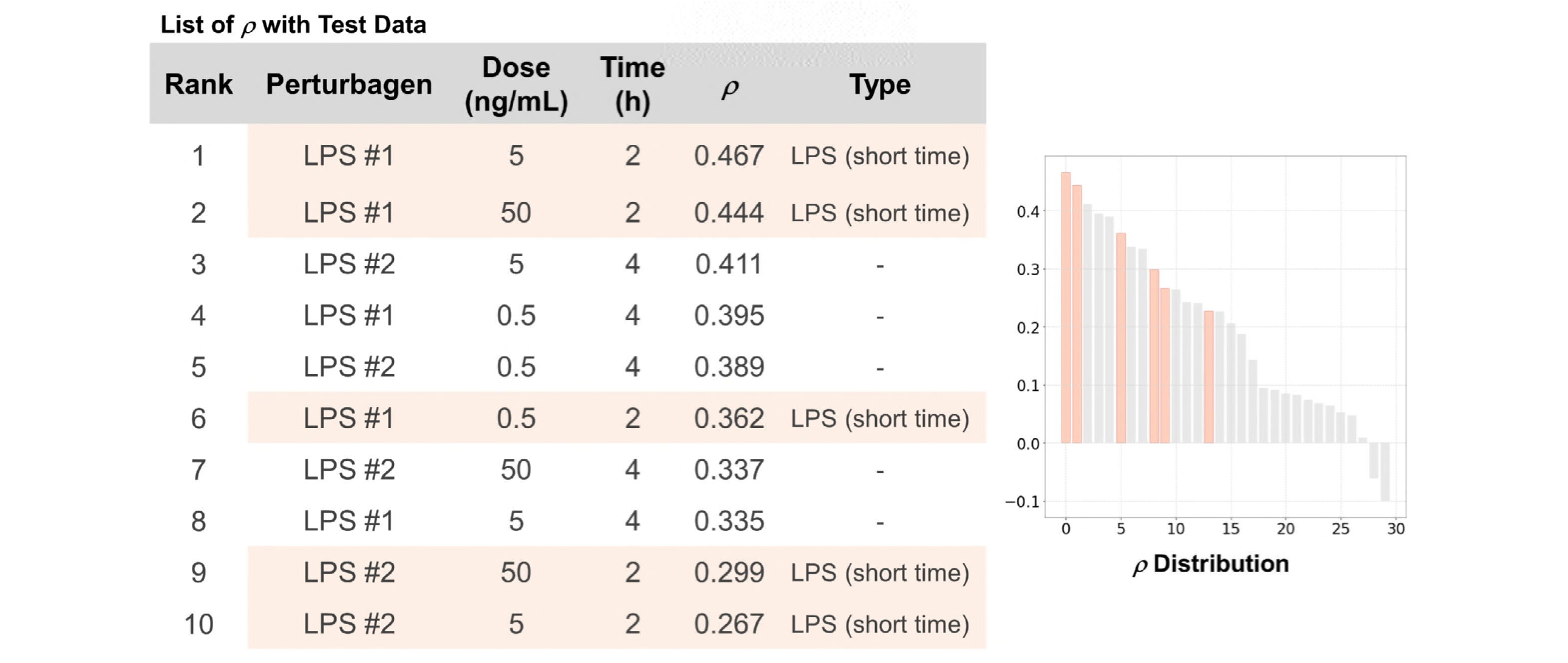
Analysis of inflammatory responses in macrophages. A. Data sets employed in this figure. B. The cumulative contribution curve of the factors comprising the training data set. The contribution of each factor to the total deviation was calculated and arranged in descending order. The cumulative contribution was calculated from the top and plotted. C. Plot of the factors whose main constituents exhibit significant enrichment of gene ontology. Genes constituting a response vector were sorted by the square of each value. The top 1% of genes were subjected to GO (biological process) analysis using Enrichment analysis of Gene Ontology Consortium. Factors annotated with significant enrichment of GO after multiple-testing corrections (Benjamini-Hochberg method, α < 0.05) were depicted in yellow-filled squares. SEGR, significant enrichment of GO. D. Analysis of the P5 factor. P5 factor scores and rho (*ρ*) of all compounds are arranged in descending order and plotted on the “Score Distribution” graph and *“ρ* Distribution” in each data set, respectively (upper, training; lower, test). Green or light salmon in the graph indicates 24-h LPS treatment. The rank, name, dose, and score of the top 10 treatments are shown. “-”, “#”, and 5 ng-indicate without 24-h LPS treatment, the sample number of biological replicates, and 5 ng/mL treatment, respectively. E. Analysis of the P6 factor using the method described in D. Green or light salmon in the graph indicates 2-h LPS treatment. The rank, name, dose, and score of the top 10 treatments are shown. “-” indicates not 2-h LPS treatment.

LPS, a well-known endotoxin, exhibits various properties as an inflammatory stimulant by binding to Toll-like receptor 4 and the effect varies from one hour to another in macrophages (Morrison Claudia R Amura *et al*, 1998). Both replicates of LPS-24-h and 2-h treatment were ranked in the top 2 of the perturbagens list sorted by the P5 and P6 factors, respectively, and the conservation of the gene patterns in another data set was confirmed by calculating Spearman’s correlation (**Fig 4D and E**). Of note, the scores of the two factors exhibited clear inverse correlation with regard to time points (Fig EV3A and B), which supports that OLSA succeeded in extracting time-dependent responses of LPS as reported (Morrison Claudia R Amura *et al*, 1998; Correa *et al*, 2013). It should be noted that an hour LPS treatment did not correlate with the P6 factor in the test data set although the treatment was “short.” One explanation is that an hour is too short to activate the transcriptional network constituting the P6 factor.

The responses to interferon β and γ treatment for 24 h seemed to be included in the P10 and P15 factors, respectively, although we were not able to validate the responses in another data set because of a lack of data (**Fig EV3B and C**). These results indicate that OLSA works in the analysis of a response-profile matrix composed of relatively small, 30 transcriptome, data.

#### 2. Application of OLSA in understanding the effects of drugs Decomposition of Hsp90-inhibitor effect

Next, we investigated whether OLSA contributes to an understanding of the effects of drugs by analyzing the CMap data set.

Hsp90 inhibitors are potent anticancer reagents in development and their mechanism is literally the inhibition of Hsp90 (Mellatyar *et al*, 2018). The first compound identified in this class of inhibitor is geldanamycin, as found in *Streptomyces hygroscopicus* (Mellatyar *et al*, 2018). Via a lead optimization process to reduce toxicity and enhance water solubility among other things, tanespimycin and alvespimycin were synthesized, and both are in clinical trials for several types of cancer (Solarova *et al*, 2015). Monorden is also an Hsp90 inhibitor. However, it was found from *Pochonia chlamydosporia* and is structurally distinct from geldanamycin (**Fig 5A**) (Solarova *et al*, 2015).

**Figure 5.**
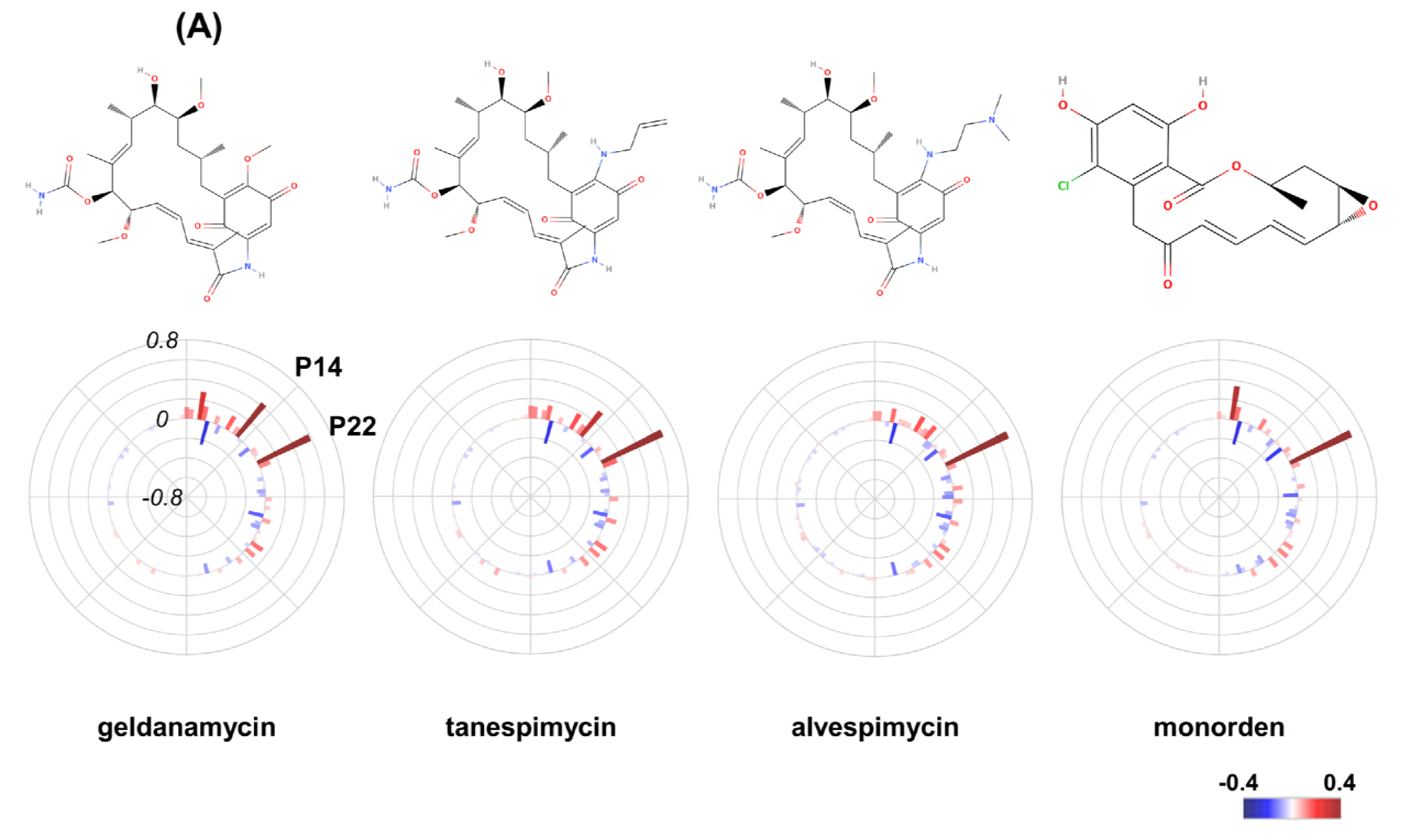
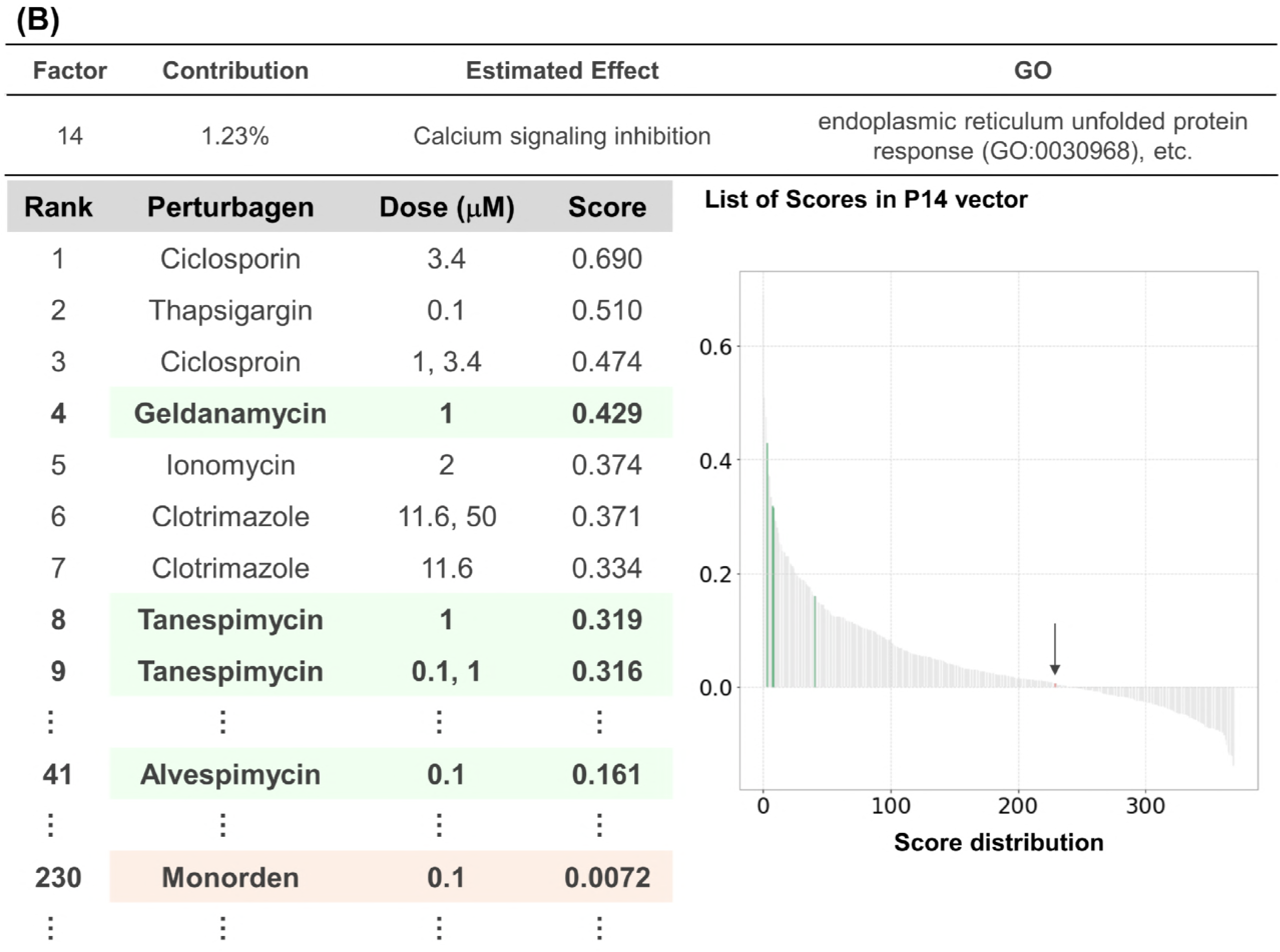
Decomposition of Hsp90-inhibitor effect. A. Structures and response scores of Hsp90 inhibitors. Structures were obtained from MolView (http://molview.org/). Response scores are plotted as a bar chart in polar coordinates with heatmap. B. Analysis of the P14 factor. P14 factor scores of all compounds are arranged in descending order and plotted on the “Score Distribution” graph. Green and light salmon with an arrow in the graph indicate geldanamycin-type inhibitors and monorden, respectively. The rank, name, dose, and score are shown.

In the OLSA result, these Hsp90 inhibitors (geldanamycin, tanespimycin, alvespimycin, and monorden) exhibited high scores in the P22 factor, which suggests that this factor includes an Hsp90 inhibition effect (**Fig EV4A**). Interestingly, there exists a difference among them as well: geldanamycin and tanespimycin exhibited high scores in the P14 factor while those of alvespimycin and monorden were not high and almost zero, respectively (**Fig 5B and Fig EV4B**). The compounds that ranked high in the P14 factor score list were cyclosporin (a calcineurin inhibitor), thapsigargin (an ER calcium depleter), ionomycin (a calcium ionophore), and they indicate the factor includes calcium signaling inhibition. Therefore, based on the P14 score, geldanamycin and tanespimycin are considered to have a high inhibitory effect of calcium signaling while the effect of alvespimycin and monorden is predicted to be mild and low, respectively. Indeed, Chang *et al* elucidated the difference between geldanamycin and monorden and reported that only the former possesses the calcium depletion effect (Chang *et al*, 2006). Moreover, alvespimycin is of lower toxicity than its lead compounds, geldanamycin and tanespimycin (Mellatyar *et al*, 2018). They are consistent with the above consideration based on the P14 factor score. Of note, all four Hsp90 inhibitors are located in quite near positions by clustering analysis, which supports the utility of OLSA in understanding Hsp90 inhibitor characteristics (**Fig EV4C**).

### Decomposition of topoisomerase-inhibitor effect

Topoisomerase inhibitors have been employed as anticancer drugs for a long time and are highly active against many types of neoplastic diseases (Delgado *et al*, 2018). Anticancer compounds, particularly anthracyclines among them, often exhibit cardiotoxicity, which restricts the application of that type of antineoplastic agent. Although it has been studied widely, the mechanism has not been clearly determined (Yu *et al*, 2018).

The OLSA results of anthracyclines (doxorubicin (also called Adriamycin), daunomycin, and mitoxantrone) revealed that the P5, P15, P16, and P17 factor scores were commonly high in topoisomerase inhibitors including nondrug compounds and the P17 factor stood out among them (**Fig EV5**). In addition to topoisomerase inhibitors, GW-8510 (a CDK2 inhibitor) and staurosporine (a multiple kinase inhibitor) were included in the high score compounds in the P17 factor. Therefore, the P17 factor is estimated to be one of the main effects of topoisomerase inhibitors and includes G1/S arrest (Shapiro, 2006; Morrison Claudia R Amura *et al*, 1998; Murray *et al*, 2013). Indeed, H-7 (a multiple kinase inhibitor with topoisomerase inhibition activity), GW-8510, and alsterpaullone (a multiple CDK inhibitor) exhibited high Spearman correlation coefficients with the P17 factor in the test PC3 data set (**Fig EV5A**). It is consistent with the above estimation because there is no other reported topoisomerase or CDK inhibitor in the test data set.

The P15 and P16 factor constituting genes were subjected to GO analysis and the genes termed “mitochondrial ATP synthesis coupled electron transport (G0:0042775)” and “organelle organization (G0:0006996)” were respectively enriched in the gene sets (**Fig EV5B**). However, we were not able to detect the commonality of the compounds in those factors other than topoisomerase inhibitors and not able clearly to determine the cellular responses of the factors although P15 constituting genes seem to be associated with mitochondria (**Fig EV5C and D**). By contrast, the P5 factor was annotated with Na+/K+ ATPase inhibition in **Figure 2**. Several studies reported Na+/K+ ATPase inhibition by doxorubicin (Gosã *et al*, 1979). Notably, one of the mechanisms explaining the cardiotoxicity induced with topoisomerase inhibitors is the inhibition of Na+/K+ ATPase (Solomonson & Halabrin, 1981). This hypothesis is consistent with the relatively high scores for the P5 factor that topoisomerase inhibitors exhibited, although the cardiotoxicity-causing mechanism remains to be determined (**Fig 5B**).

These results indicate that it is possible to decompose the effect of a drug with OLSA and imply that OLSA can detect not only the main effect of a drug but also other effects that may be the cause of toxicity or adverse events.

### Identification of autophagy regulators

Finally, we explored the possibility of OLSA for drug repositioning. The analysis of CMap-derived data suggested that the P2 factor includes a basic effect responding to PI3K/AKT/mTOR signaling inhibition (**Fig EV6A**). Mammalian target of rapamycin complex (mTORC) I is a critical regulator of autophagy and its inhibition affects many essential cellular phenomena (Dibble & Cantley, 2015). In the literature, we noticed that many compounds in the top 10% (37) of the list sorted by P2 factor scores were reported to be autophagy inducers although LY-294002 and wortmannin are not autophagy inducers because they have two-sided effects because of subclasses of PI3K (Table 1) (Wang *et al*, 2015). By contrast, there was no information regarding any autophagy relationship in 8 compounds on the list: 0297317-0002B, thonzonium bromide, benzethonium chloride (BC), methylbenzethonium chloride, phenazopyridine (PP), benzamil (BEN), metiothepine (MTP), and metixene (MTX). Therefore, we hypothesized that those compounds were associated with autophagy regulation and tested the hypothesis. Among them, 0297417-0002B, tonzonium bromide, and methylbenzethonium chloride were excluded from the test compounds because the former was not easily available and the latter two were the same type of cationic detergent as BC. The remaining compounds, BC, BEN, MTP, MEX, and PP were subjected to *in vitro* analysis (**Fig 7A, B, and C**). Interestingly, those 5 compounds were not clustered with typical autophagy regulators such as sirolimus (also called rapamycin) and wortmannin by a hierarchical clustering method (**Fig EV6B**). HeLa cells, a well-used human cervical cancer-derived cell line, were treated with the tested compounds and loperamide (LOP), as a positive control (Kaizuka *et al*, 2016) and the conversion of LC3-I to LC3-II was examined with western blotting analysis. We employed LOP because it exhibited similar scores to those tested compounds and was reported as an autophagy inducer in more than two independent reports (Zhang *et al*, 2007; Kaizuka *et al*, 2016). The conversion, one of the indicators of autophagy induction (Klionsky *et al*, 2016), was clearly increased by 4 of 5 compounds (BC, BEN, MTP, and MTX) while PP had almost no effect (**Fig 7D**).

**Figure 6.**
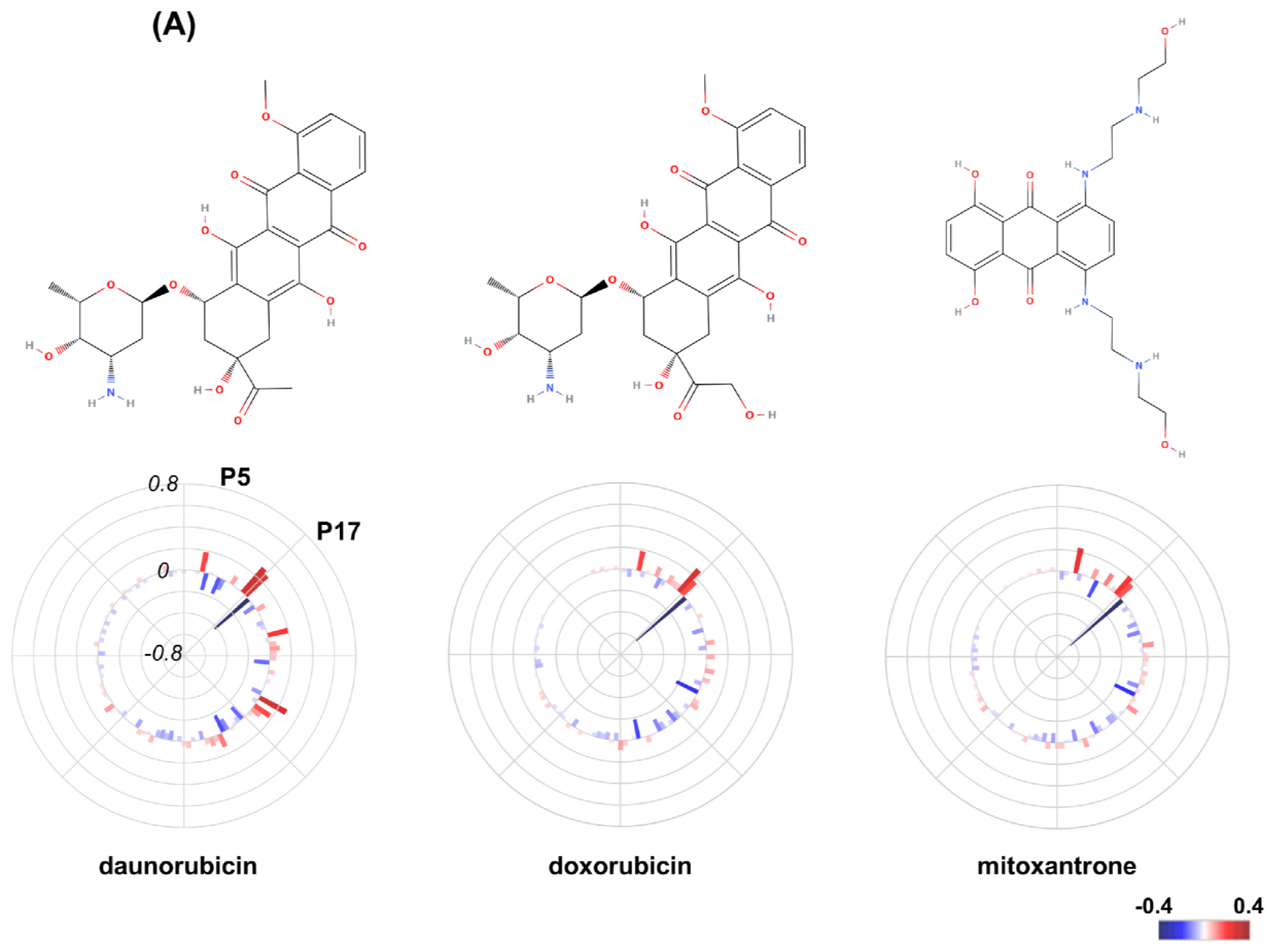
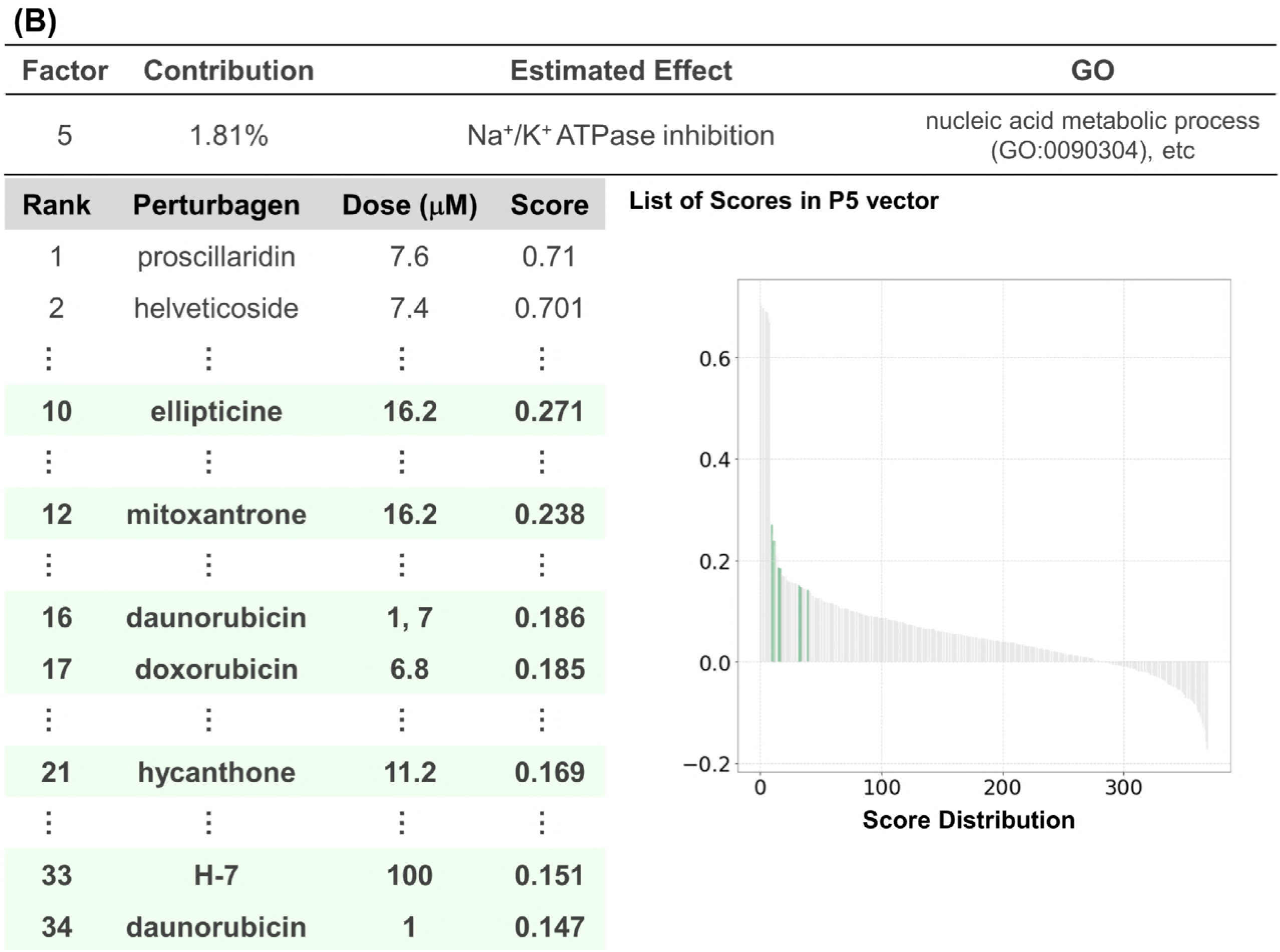
Decomposition of topoisomerase-inhibitor effect. A. Structures and polar charts of response scores of topoisomerase inhibitors: daunorubicin, doxorubicin, and mitoxantrone. For daunorubicin, 7 μM dose data were employed considering the higher effect on the transcriptional network than that of 1 μM. Structures were obtained from MolView (http://molview.org/). Response scores are plotted as a bar chart in polar coordinates with heatmap. B. Analysis of the P5 factor. P5 factor scores of all compounds are arranged in descending order and plotted on the “Score Distribution” graph. Green in the graph indicates topoisomerase inhibitors. The rank, name, dose, and score are shown.

**Figure 7.**
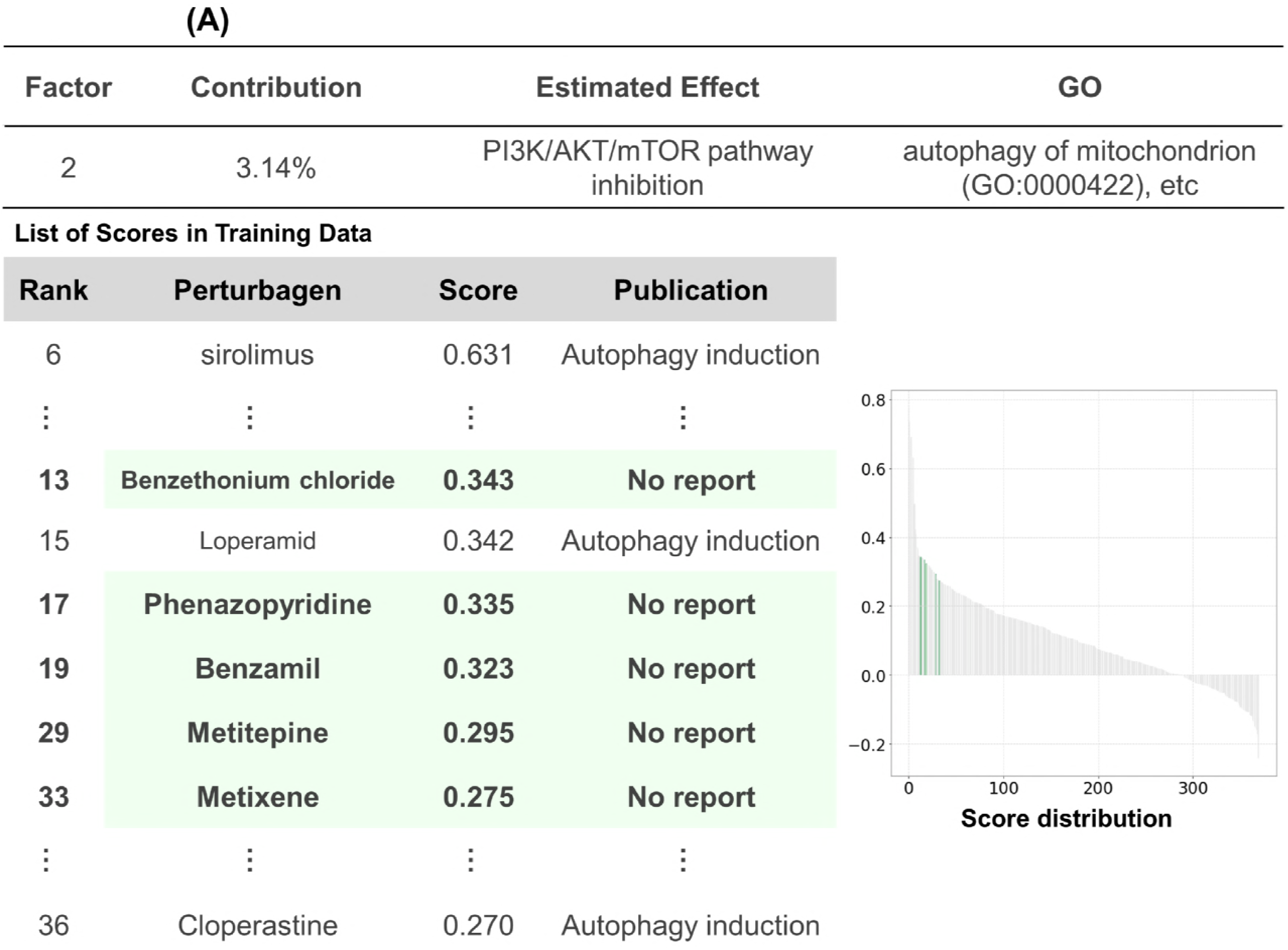
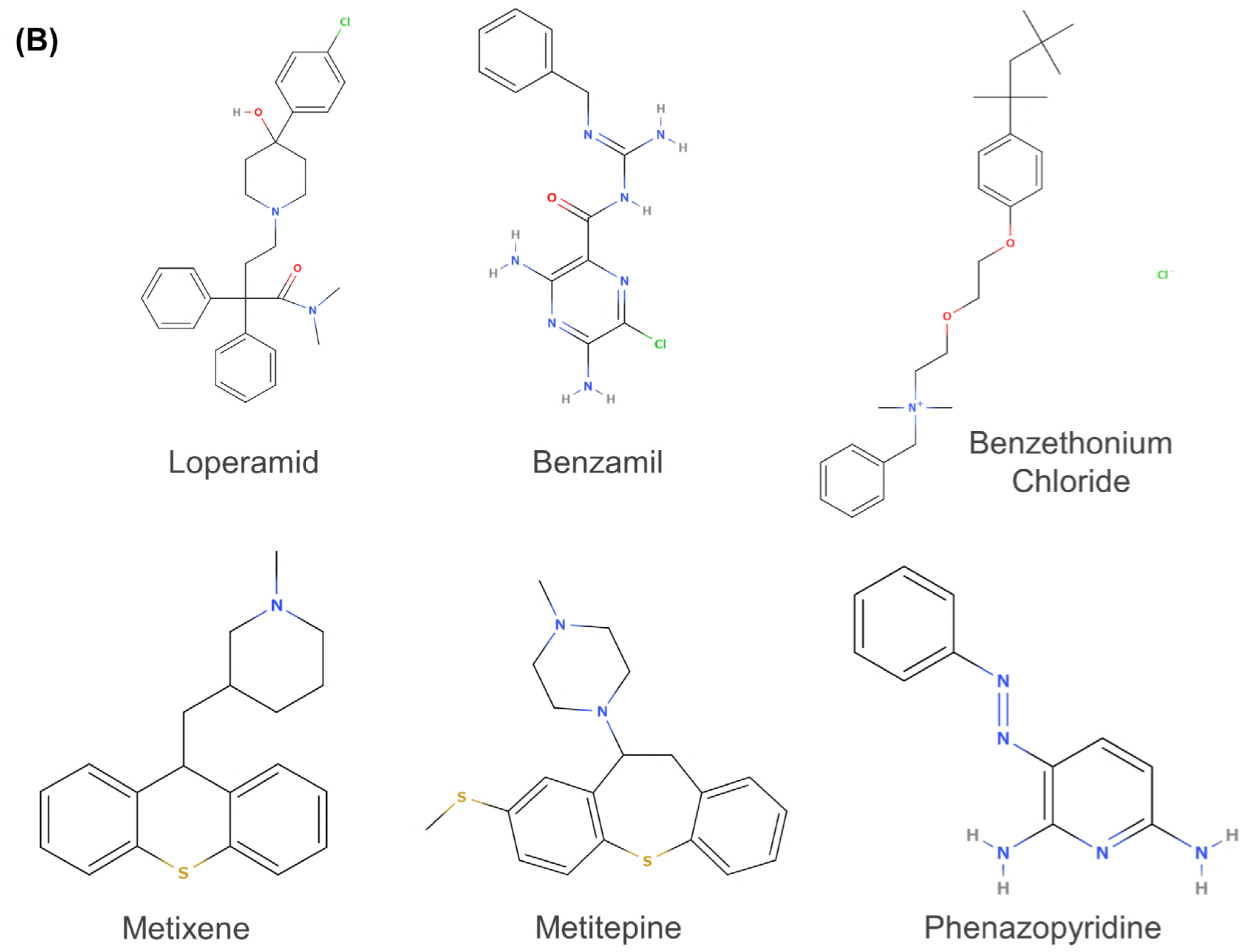
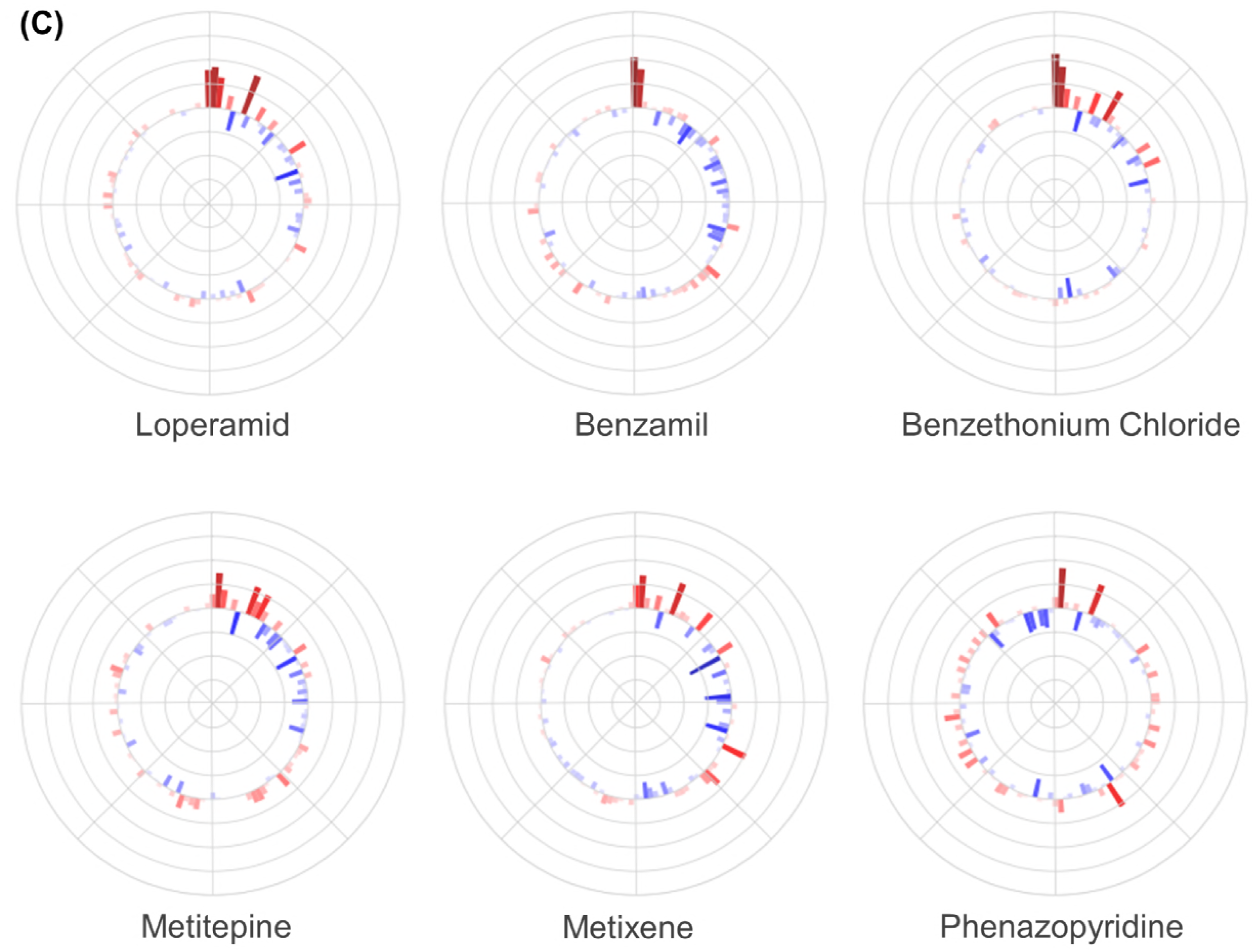
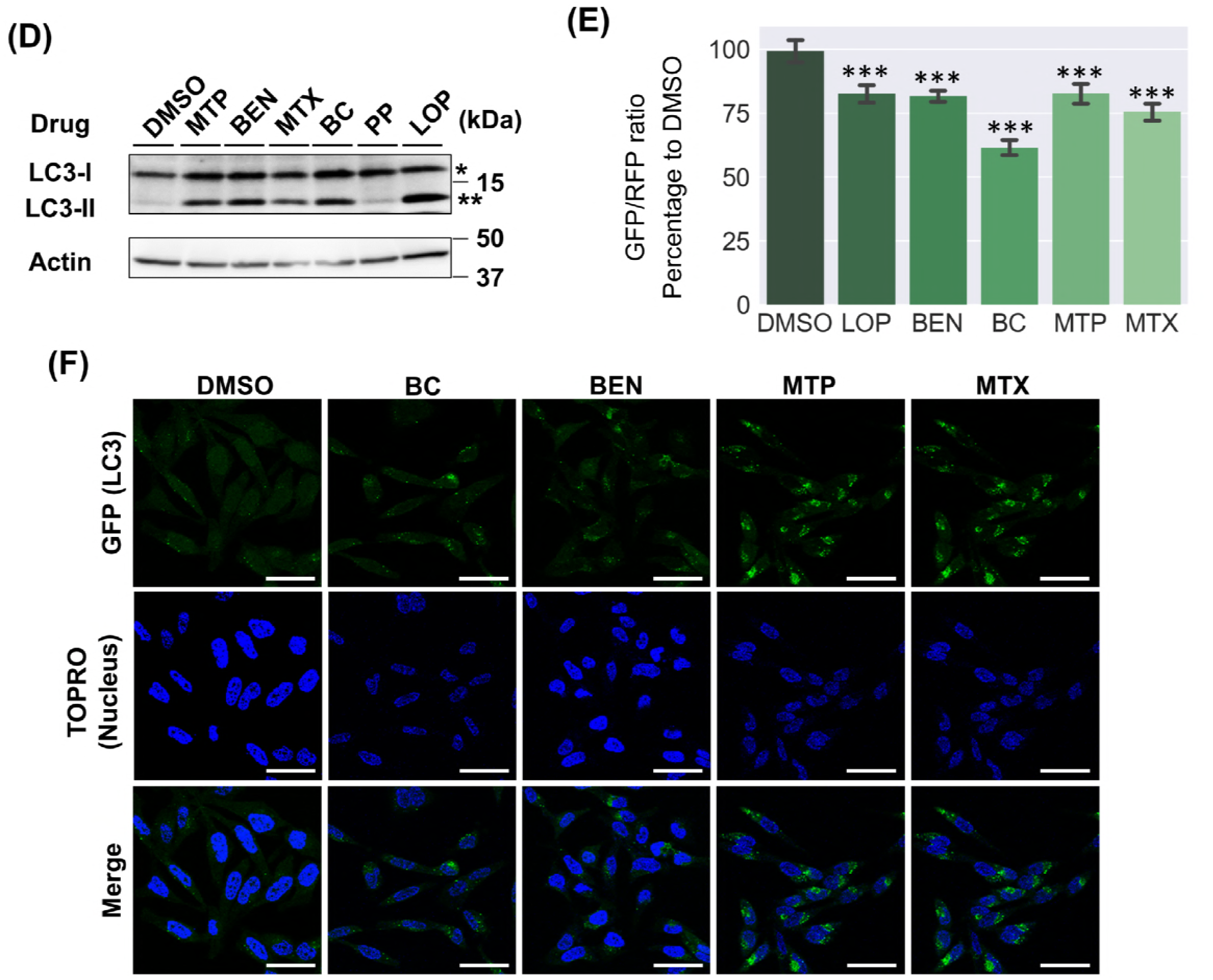
Identification of autophagy regulators. A. Analysis of the P2 factor. P2 factor scores of all compounds are arranged in descending order and plotted on the “Score Distribution” graph. Green in the graph indicates the compounds with a high P2 score but without reports about autophagy. The rank, name, dose, and score are shown. B. Structures of the tested compounds in this study. Structures were obtained from MolView (http://molview.org/). C. Polar charts of response scores of the tested compounds. Response scores are plotted as a bar chart in polar coordinates with heatmap. D. Western blotting analysis of HeLa cells treated with the tested compounds. HeLa cells were treated with the tested compounds at the indicated concentration for 24 h. The whole-cell lysate was analyzed by western blotting using anti-LC3 antibody. *LC3-I, **LC3-II. E. Autophagy flux evaluation of GFP-LC3-RFP-LC3ΔG-HeLa cells treated with the tested compounds. HeLa cells expressing GFP-LC3-RFP-LC3ΔG were treated with the tested compounds using the method described in D. GFP and RFP signals were quantified with a Tecan Infinite M200 plate reader and the GFP/RFP ratio was calculated. Each bar represents the mean ± SE, *n* = 6. Significance test was conducted with Turkey-Kramer method and significant differences between DMSO and the tested compounds are only shown: ***P < 0.001. F. Imaging analysis of GFP-LC3-RFP-LC3ΔG-HeLa cells treated with the tested compounds. HeLa cells expressing GFP-LC3-RFP-LC3ΔG were treated with the tested compounds using the method described in D, fixed with 4% paraformaldehyde, stained with TO-Pro-3 iodide, and the fluorescence signals were detected with a TCS SP5 confocal microscope. Green and blue signals indicate GFP (LC3) and TO-Pro-3 iodide (nucleus), respectively. Scale bars correspond to 50 μm. In Fig 7D, E, and F, a representative result of at least two independent experiments is shown.

**Table 1.** List of P2 factor high score compounds in the MCF7 cells’ data set of CMap. Rank, name, concentration, score, and reference are shown in this table.

| Rank | Compound | Score | Dose ($\mu\text{M}$) | Reference |
| --- | --- | --- | --- | --- |
| 1 | wortmannin | 0.776 | 0.01, 1 | (Hansen *et al*, 1995) |
| 2 | wortmannin | 0.735 | 0.01 | (Hansen *et al*, 1995) |
| 3 | LY-294002 | 0.691 | 10 | (Vlahos *et al*, 1994) |
| 4 | LY-294002 | 0.69 | 0.1, 10 | (Vlahos *et al*, 1994) |
| 5 | wortmannin | 0.632 | 1 | (Hansen *et al*, 1995) |
| 6 | sirolimus | 0.631 | 0.1 | (Wu *et al*, 2013) |
| 7 | quinostatin | 0.497 | 10 | (Yang *et al*, 2007) |
| 8 | trifluoperazine | 0.423 | 8.4 | (Zhang *et al*, 2007) |
| 9 | 0297417-0002B | 0.408 | 10 | No report |
| 10 | trifluoperazine | 0.371 | 8.4, 10 | (Zhang *et al*, 2007) |
| 11 | tetrandrine | 0.367 | 6.4 | (Wong *et al*, 2017) |
| 12 | tonzonium bromide | 0.347 | 6.8 | No report |
| **13** | **benzethonium chloride** | **0.343** | **9** | **No report** |
| 14 | methylbenzethonium chloride | 0.343 | 8.6 | No report |
| 15 | loperamide | 0.342 | 7.8 | (Zhang *et al*, 2007) |
| 16 | troglitazone | 0.34 | 10 | (Wu *et al*, 2013) |
| **17** | **phenazopyridine** | **0.335** | **16** | **No report** |
| 18 | prochlorperazine | 0.325 | 6.6, 10 | (Cheng *et al*, 2015) |
| **19** | **benzamil** | **0.323** | **11.2** | **No report** |
| 20 | prochlorperazine | 0.321 | 10 | (Cheng *et al*, 2015) |
| 21 | fluphenazine | 0.319 | 10 | (Barmada *et al*, 2014) |
| 22 | thioridazine | 0.317 | 10 | (Cheng *et al*, 2015) |
| 23 | fluphenazine | 0.317 | 7.8, 10 | (Barmada *et al*, 2014) |
| 24 | resveratrol | 0.311 | 10 | (Wang *et al*, 2017) |
| 25 | trifluoperazine | 0.306 | 10 | (Zhang *et al*, 2007) |
| 26 | thioridazine | 0.305 | 9.8 | (Cheng *et al*, 2015) |
| 27 | homochlorcyclizine | 0.3 | 10.4 | (Kaizuka *et al*, 2016) |
| 28 | perhexiline | 0.298 | 10.2 | (Balgı *et al*, 2009) |
| **29** | **metiothepine** | **0.295** | **8.4** | **No report** |
| 30 | thioridazine | 0.289 | 0, 1, 10 | (Cheng *et al*, 2015) |
| 31 | fulvestrant | 0.278 | 0.01, 1 | (Yu *et al*, 2016) |
| 32 | amiodarone | 0.278 | 5.8 | (Balgı *et al*, 2009) |
| **33** | **metixene** | **0.275** | **11.6** | **No report** |
| 34 | maprotiline | 0.275 | 12.8 | (Cloonan & Williams, 2011) |
| 35 | raloxifene | 0.273 | 7.8 | (Kim *et al*, 2015) |
| 36 | cloperastine | 0.27 | 11 | (Kaizuka *et al*, 2016) |
| 37 | fulvestrant | 0.267 | 1 | (Yu *et al*, 2016) |

Next, we employed a new autophagic flux probe, GFP-LC3-RFP-LC3ΔG, developed by Dr. Kaizuka *et al* to investigate the capacity of the 4 positive compounds precisely as autophagy regulators. Endogenous ATG4 protease cleaves the probe into equimolar amounts of GFP-LC3 and RFP-LC3ΔG. The former is degraded in lysosome via autophagosome while the latter remains in the cytosol and works as an internal control. Thus, calculation of the GFP/RFP fluorescence ratio enables the precise estimation of autophagic flux compared with WB analysis of LC3-I to LC3-II conversion (Morishita *et al*, 2017). GFP-LC3-RFP-LC3ΔG-expressing HeLa cells were treated with BC, BEN, MTP, and MTX and the GFP/RFP fluorescence ratio was measured. As reported, LOP (an autophagy inducer) and bafilomycin A1 (an autophagy inhibitor) decreased and increased the signal ratio, respectively, and all 4 compounds reduced the ratio, which suggests they are autophagy inducers (**Fig 7E**). GFP-LC3 puncta were clearly observed under the same condition (**Fig 7F**). All of the results indicate that BC, BEN, MTP, and MTX are autophagy inducers. Four of five compounds predicted to be autophagy regulators by OLSA actually induced autophagy, which supports one of the utilities of this method in drug repositioning or rescue.

## Discussion

Drug discovery with drug repositioning has been a successful approach (Tan *et al*, 2018; Mercorelli *et al*, 2018; Panchapakesan & Pollock, 2018). The approach mostly depends on serendipity in the beginning, but the various methodologies based on scientific evidence such as screening systems and *in silico* prediction of drug–molecule interaction have been established, which contribute to the development of the approach (Bellomo *et al*, 2017; Datta *et al*, 2018; Karaman B, 2018). The success of drug repositioning implies that the effect of a drug is complex and composed of multiple more basic components. Therefore, in the present study, we attempted to separate the complex effect of a drug into basic effects to understand the pharmacological properties of a drug. To achieve separation, we focused on profile data analysis. The important characteristic of omics is the conversion of the biological information of a sample into numeric values by its comprehensiveness, which enables mathematical approaches to the analysis of biological samples. However, in general, raw values of omics data are multiple variables, complex and often difficult to comprehend because of the curse of dimensionality (Adachi, 2017). Analytical methods setting a new layer that appropriately contracts the degrees of freedom are indispensable for extracting information from the data and many methods have been devised (Aziz *et al*, 2017; Ren *et al*, 2013). The factors generated with OLSA are expected to be the exact new indicators constituting a novel layer and, therefore, we can regard OLSA as one of the analytical methods to set a new layer compressing biological information.

We should estimate the biological meanings of the new indicators cautiously because the contraction of profile data with OLSA is conducted in an unsupervised way and the interpretation depends on the analysts. We have tried to estimate using two approaches: the first is to analyze the variables mainly constituting a factor and another is to utilize the similarity of the high-ranked samples in the list sorted by the response-score ranking of the focusing factor. When transcriptome profile data were subjected to OLSA, the variables were genes. There are well-established methods of analysis of the relationship of genes, such as GO analysis and pathway analysis (Ashburner *et al*, 2000; Garcia-Campos *et al*, 2015). To obtain insight about whether a factor has consistent biological meanings or not, we employed statistical significance in GO analysis of the main genes constituting the focused factor and evaluated the correspondence to the existing bodies of knowledge as a requirement. Interestingly, the ratio of the factors with significant enrichment of GO varied between the data sets and SEGR was 0.551, 0.724, and 0.333 in CMap, HepG2, and BMDM response-profile matrices, respectively (**Fig 2C, Fig 3C, and Fig 4C**). An explanation of the differences is the possibility that the contribution ratio of biologically relevant factors is different between the data sets. In the data sets with relatively low SEGR (CMap and BMDM set), the factors annotated with GO tended to exhibit high contribution to the total deviation and the significance of the enrichment was supported by the results of Fisher’s exact tests (**Fig EV7**). Moreover, in some of the factors with a high contribution ratio, but without GO annotation, for instance the P26 and P35 factors, structurally similar compounds dominate the top of the score ranking, which implies the association with some cellular responses (**Fig 2 and EV1C**). OLSA is a matrix-decomposition method based on PCA and a factor with a high contribution ratio means that the factor includes a response with large variance. As GO is a classification method based on existing knowledge, the relation between factor contribution and annotation with GO in our results is consistent with our daily experience: a phenomenon with large phenotype is easily detected while a faint phenotype is often missed.

What is the difference between the biologically relevant factors and those that are not? We consider that the difference is whether the contracted deviation is derived from random noise or not. In general, error is divided into two types, systematic and random. Systematic error can be regarded as a response of cells to perturbation and may be separated as a factor although the perturbagen may not be physiologically meaningful, such as deviation in experimental techniques or batch differences. The latter is important here because the random error is derived from technical restraints such as the limit of quantitation and these are not biologically relevant. This is consistent with the difference of the factor correspondence to existing knowledge, GO, between the data sets as quantitation of genes depends on system limitations such as those in assay systems or sample conditions, and varies between experiments.

The biologically relevant factors can be classified into 3 types: (1) well characterized, (2) not characterized or identified, and (3) nonlinear and cannot be separated. Because OLSA is an unsupervised analysis, it is difficult to distinguish between noise, uncharacterized, and the unseparated factors. However, mathematical considerations focusing on the contribution ratio may be useful for setting the criteria, although we have applied the generally used threshold in PCA and analyzed the cumulative contribution ratio of factors in the present study (Lever *et al*, 2017). The relationship between factor contribution and biological relevance is an important point of OLSA and its analysis is an essential future task.

We consider that OLSA is a research tool for assessing the purity of the effects of a group of candidate compounds and thereby facilitating the lead optimization process for drug discovery (Roy, 2018). OLSA provides scores of the common factors in the group and compound-specific factors among cellular responses to the candidate compounds. Because the former is considered to include the effects based on the MOA of the candidates—so-called “class effects”—we can prioritize the candidates according to the purity of their effects based on the scores of the common effects, leading to a deeper understanding of their structure–activity relationships, and the rational design of potential drugs. Identifying the purity of the effects of candidates is generally expected to be useful for avoiding toxicity specific to a candidate, which is often difficult to determine, whereas compound-specific effects can contribute to the therapeutic effectiveness of the selected candidate for the target disease. For example, in the analysis of topoisomerase inhibitors in the CMap data set, the scores of those inhibitors were commonly high in the P15 factor, which was annotated with mitochondria toxicity in GO analysis. This is consistent with several studies that reported mitochondrial toxicity of topoisomerase inhibitors (Filipa *et al*, 2014), which could contribute to the mechanism of their antitumor activity. Thus, OLSA is expected to be useful for determining the biochemical assays necessary for the next step in the lead optimization process of drug discovery.

In the present study, we tested whether the factors obtained by OLSA could be utilized for drug repositioning, focusing on the P2 factor in CMap data; a factor expected to include PI3K/AKT/mTOR inhibition and to be associated with autophagy via mTORC1 modulation. Western blotting and the following analysis using a novel autophagic flux probe system revealed that 4 of 5 tested compounds actually induced autophagy, while this prediction was not achieved by conventional clustering analysis (**Fig EV6B**). The results support the potency of the strategy of drug effect separation. However, we should carefully discuss why PP did not induce autophagy, contrary to the prediction. One of the reasons is considered to be the discrepancy between the wanted cellular response for repositioning and the estimated response, particularly in cases where the cellular response is regulated by several factors. For instance, LY294002 and wortmannin are listed as P2V high score compounds and reported to inhibit mTORC1 activity as predicted, but do not induce autophagy because they are pan-inhibitors of PI3K (Takeuchi *et al*, 2005; Wu *et al*, 2010). The PI3K family is divided into three classes (Wang *et al*, 2015). Class I PI3K activates mTOR via PI3K/AKT/mTOR signaling and reduces autophagy, while Class III PI3K increases phagophore formation and promotes autophagy. Because of this two-sided effect, pan-PI3K inhibitors do inhibit mTORC1 but do not induce autophagy (Takeuchi *et al*, 2005; Wu *et al*, 2010). We confirmed that PP treatment decreased phosphorylation of S6K, an mTORC1 substrate and an mTORC1 activation marker, which indicates that PP does have an mTORC1 inhibitory effect as predicted (**Fig EV5**). Thus, OLSA actually contributed to the understanding of a perturbagen effect by separating it, whereas the wanted cellular response for drug repositioning does not always correspond directly to the estimated response. It may be helpful for drug repositioning strategy with OLSA to consider the relationship among the factors using techniques including graphical modeling (Djordjilović *et al*, 2015).

OLSA is applicable not only to transcriptome data but also to the other layers of omics data. Considering the central dogma, data in the proteome or metabolome may be more suitable for the analysis to understand the effects of drugs on cells. Linearity is essential in OLSA and is considered to depend on the characteristics of each layer. A comparison of the OLSA results for many drugs among the layers such as transomics is informative and will contribute to an understanding of the characteristics of each layer, and to determine which layer is suitable for evaluating a particular type of drug. Moreover, OLSA does not require existing knowledge or biological meaning of the variables in profiles, and thus, can be utilized for analyzing the data composed of the variables without biological information, such as the spots in two-dimensional electrophoresis and features in phenotyping screening (Muroi *et al*, 2010). Although the databases summarizing such omics data are not substantial today and we were not able to test them in the present study, this challenge is important.

In the present study, we proposed the potential of OLSA in the separation of a drug effect and the extraction of more basic components. We expect that this method and further analyses will contribute to drug repositioning, lead optimization, and other approaches in drug discovery.

## Materials and Methods

### Data preprocessing for OLSA input

The expression data matrix was prepared as variables in rows and samples in columns. Here, we describe the procedures considering transcriptome data although the methodology can be applied to other types of omics data. Data from each sample was converted into a nonparametric rank-ordered list of all genes in the transcriptome data based on the expression values (expression rank matrix). To obtain differential expression values to the controls, we employed a robust z-scoring procedure and the differential expression value gene x to the control in the ith sample was computed as:

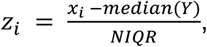

where x and *Y* are the vectors of gene x ranks across all samples and the control samples in the expression rank matrix, respectively, and *NIQR* is the normalized interquartile range of the control sample values. We call the obtained matrix a response-profile matrix.

### OLSA

OLSA consists of the following procedures.

#### 1. Data selection

To exclude the samples that are considered to lose reversibility, L2-norm of each sample is calculated, and the outlier samples determined by the Smirnov-Grubbs test are removed from the response-profile matrix. We call the diagonal matrix consisting of L2-norm “total strength.”

#### 2. Normalization

The selected data are then normalized by L2-norm.

#### 3. Mirror data preparation

A point-symmetric data set to the normalized data set is generated and combined with the original data set, which makes the centroid of the combined data set zero.

#### 4. Principal component analysis

The generated data set is subjected to principal component analysis to contract the gene expression changes with their coordination.

#### 5. Varimax rotation

To reduce the genes contributing to each contracting vector retaining the orthogonality of the vectors, the varimax rotation is applied to the obtained matrix consisting of the contracting vectors. In this paper, the vectors are sorted by contribution ratio and the ones from the top to the vector whose cumulative contribution ratio exceeds 0.8 (CMap and Magkoufopoulou’s data) or 0.9 (Raza’s data) are subjected to varimax rotation considering calculation time. We define the rotated vectors as “response vectors” or “factors” and call a matrix consisting of response vectors the “response-vector matrix.”

#### 6. Generation of response-score matrix

Employing the response-vector matrix, the scores of the samples for each factor are calculated as inner products. We label a matrix consisting of the scores the “response-score matrix.”

### GO analysis of genes mainly constituting the response vector

Genes constituting a response vector were sorted by the square of each value. The top 1% of genes were subjected to GO (biological process) analysis using Enrichment analysis of Gene Ontology Consortium (http://www.geneontology.org/). The obtained p-values were processed using the Benjamini-Hochberg method for multiple-testing corrections among factors (α < 0.05).

### Validation of biological meaning of response vector

The validation of the estimated biological response of a response vector annotated with GO, the treatment similarity, or the results of a literature survey was conducted by investigating Spearman’s correlations between the factor constituents and the samples in another data set. First, the contribution of each gene constituting a response vector was calculated as the ratio of the square of each gene value to the summation of those values and then sorted by the ratio. Genes from the top to the gene whose cumulative contribution ratio exceeded 0.9 in the list were selected and employed as the signature representing the factor for calculation of Spearman’s correlation.

### Materials

Benzethonium chloride (025-11662), loperamide hydrochloride (129-05721), and phenazopyridine hydrochloride (162-14441) were purchased from Wako Pure Chemical Industries (Osaka, Japan). Methiothepine hydrochloride (sc-253005) and anti-β-Actin (sc-47778) were purchased from Santa Cruz Biotechnology (Dallas, TX). Benzamil (3380) was purchased from Tocris Bioscience (Bristol, UK). Metixene hydrochloride (M1808000) and bafilomycin A1 from *Streptomyces griseus* (B1793) were purchased from Sigma-Aldrich (St. Louis, MO). Rabbit anti-p70 S6 kinase (9202) and mouse anti-phospho-p70 S6 kinase (9206) were purchased from Cell Signaling Technology (Beverly, MA). Rabbit anti-LC3 (PM036) was purchased from Medical & Biological Laboratories (Nagoya, Japan). All other chemicals were of analytical grade.

### Cell culture

GFP-LC3-RFP-LC3ΔG-expressing HeLa cells were cultured in Dulbecco’s Modified Eagle’s Medium (DMEM) (D6546, Sigma-Aldrich) supplemented with 10% fetal bovine serum (FBS) and 2 mM L-glutamine (G7513, Sigma-Aldrich). HeLa cells (CCL-2, ATCC) were cultured in DMEM (10313-021, Life Technologies, Carlsbad, CA) with 10% FBS and 1% MEM nonessential amino acids (11140-050, Life Technologies). All cells were maintained at 37°C under 5% CO2.

### Western blotting analysis

Western blotting analysis was conducted as previously described (Mizuno *et al*, 2015). Specimens were separated with SDS-PAGE on a 13.5% polyacrylamide gel with a 3.75% stacking gel at 140 V for 90 min. The molecular weight was determined using Precision Plus Protein Standards (1610373, Bio-Rad, Richmond, CA). Proteins were transferred electrophoretically to a poly(vinylidene difluoride) (PVDF) membrane (Pall, NY) using a blotter (Bio-Rad) at 100 V for 60 min. Nonspecific binding sites on the membrane were blocked with PVDF Blocking Reagent for Can Get Signal (Toyobo, Osaka, Japan) at room temperature for 60 min. After blocking, the PVDF membrane was incubated with primary antibodies diluted with Can Get Signal solution 1 (Toyobo) at 4°C for 24 h. Primary antibodies were used in the following conditions: anti-β-actin (1/2,000), anti-p70 S6 kinase (1/2,000), anti-phospho-p70 S6 kinase (1/2,000), and anti-LC3 (1/2,000). After the reaction with primary antibodies, the membrane was incubated with horseradish peroxidase-conjugated anti-rabbit or anti-mouse IgG antibody (Amersham Biosciences, Piscataway, NJ) diluted to 1/10,000 by Tris-buffered saline containing 0.05% Tween 20 at room temperature for 60 min. Immunoreactivity was detected with a Fusion Solo S (Vilber Lourmat, Marne la Vallée, France) and Westar ETA C Ultra 2.0 (Cyanagen, Bologna, Italy). The band intensity indicating each protein was quantified by Multi Gauge software (Fujifilm, Tokyo, Japan).

### Evaluation of autophagic flux with GFP-LC3-RFP-LC3ΔG-expressing HeLa cells

Autophagic flux after drug treatment was determined essentially as described previously (Kaizuka *et al*, 2016). GFP-LC3-RFP-LC3ΔG-expressing HeLa cells were seeded in black/clear bottom 96-well plates (353948, Corning, NY) at 1.5 × 10^4^ cells/well and maintained for 72 h. After drug treatment for 6 h, cells were washed with PBS (+), fixed with 4% paraformaldehyde solution (163-20145, Wako) for 10 min, and washed with PBS (+). Measurement of GFP and RFP fluorescence was performed using a microplate reader (Infinite M200 microplate reader; Tecan, Mannedorf, Switzerland) with excitation/emission at 480/510 nm and 580/610 nm, respectively.

### GFP-LC3 imaging

Fluorescence microscopy was conducted as described previously (Mizuno *et al*, 2011). GFP-LC3-RFP-LC3ΔG-expressing HeLa cells were seeded on glass coverslips (Matsunami Glass, Osaka, Japan) in 12-well plates at 1.5 × 10^4^ cells/well and maintained for 72 h. After drug treatment for 24 h, cells were washed twice with PBS, fixed with 4% paraformaldehyde solution for 10 min, permeabilized with 0.1% saponin in PBS for 10 min, and blocked with 3% bovine serum albumin (BSA) in PBS for 30 min. After blocking, cells were incubated with TO-Pro-3 iodide (Life Technologies, Carlsbad, CA) diluted to 1/2,500 by 2% BSA in PBS for 60 min. Coverslips were mounted with H-1000 Vectashield mounting medium (Vector Laboratories, Burlingame, CA) and analyzed with a TCS SP5 confocal microscope (Leica, Solms, Germany). Images were processed with LAS AF (Leica).

### Statistical analysis

A Student two-tailed unpaired *t* test and one-way analysis of variance followed by Tukey’s multiple comparison test were used to identify significant differences among groups, where appropriate. The data were analyzed using Prism software (GraphPad Software, La Jolla, CA).

### Data and software availability

The computer code produced in this study is available in Code EV1 and in the following database:

- OLSA python scripts: GitHub (https://github.com/tadahaya222/OLSApy)

Figure source data for **Fig 2, Fig 3, Fig 4, Fig 6, and Fig 7** are available in Figure Source Data corresponding to each figure.

Data references are listed below:

Lamb J, Crawford ED, Peck D, Modell JW, Blat IC, Wrobel MJ, Lerner J, Brunet JP, Subramanian A, Ross KN, Reich M, Hieronymus H, Wei G, Armstrong SA, Haggarty SJ, Clemons PA, Wei R, Carr SA, Lander ES, Golub TR (2006). The Connectivity Map: using gene-expression signatures to connect small molecules, genes, and disease. *Science* 313:1929-35.
Magkoufopoulou C (2011). *Gene Expression Omnibus* GSE28878 [DATASET] (https://www.ncbi.nlm.nih.gov/geo/query/acc.cgi?acc=GSE28878)
Raza S, Freeman TC, Hume DA (2013). *Gene Expression Omnibus* GSE44292 [DATASET] (https://www.ncbi.nlm.nih.gov/geo/query/acc.cgi?acc=GSE44292)

## Acknowledgements

We thank Professor Noboru Mizushima and Dr. Takeshi Kaizuka (Graduate School and Faculty of Medicine, the University of Tokyo) for kindly gifting GFP-LC3-RFP-LC3ΔG-HeLa cells. This study was financially supported by a Grant-in-Aid for Challenging Exploratory Research [17K19478] from the Japan Society for the Promotion of Science and Long-Range Research Initiative [17_S05-01-2] from the Japan Chemical Industry Association.

## Author contributions

TM and HK coordinated the project, designed experiments, analyzed all data, and wrote the manuscript. TM, SK, and HK conceived the concept. SK, SM, and TM contributed to the coding. TI conducted experiments. HK revised the manuscript for intellectual content. All authors approved the manuscript before submission.

## Expanded View Figure legends

### Figure EV1. Analysis of cellular responses in MCF7 cells treated with 370 perturbagens

A. Structure of flavonoids in the data sets.

B. Analysis of the P76 factor using the method shown in Fig 2D.

### Figure EV2. Analysis of cellular responses in HepG2 cells treated with 62 genotoxic compounds

A. Analysis of the P15 factor using the method described in Fig 3D. Green or light salmon in the graph indicates dimethyl nitrosamine or phenacetin. NPD, 4-nitro-*o*-phenylenediamine.

B. Analysis of the P29 factor using the method described in Fig 3D. Green or light salmon in the graph indicates reserpine or TCDD. TCDD, 2,3,7,8-tetrachlorodibenzo-*p*-dioxin; MOCA, 4,4’-methylene-bis(2-chloroaniline).

### Figure EV3. Analysis of inflammatory responses in macrophages

A. Heatmap comparing the scores of P5 and P6 factors. 1 h, &, 24 h and #1, #2 indicate 1 h-, &, 24 h treatment and the number of biological replicates, respectively.

B. Scatter plot of the scores of P5 and P6 factors. The blue line and area indicate the regression line and the 95% confidence interval. *R^2^*, the coefficient of determination.

C. Analysis of the P10 factor using the method shown in Fig 4D. Green in the graph indicates 24- or 8-h interferon β treatment.

D. Analysis of the P15 factor using the method described in Fig 4D. Green in the graph indicates 24- or 8-h interferon γ treatment.

### Figure EV4. Decomposition of Hsp90-inhibitor effect

A. Analysis of the P22 factor using the method shown in Fig 2D. Green or light salmon indicates Hsp90 inhibitors.

B. Analysis of P14 factor using the method shown in Fig 2D. Green and light salmon indicate calcium signaling modulators and Hsp90 inhibitors, respectively.

C. Clustering analysis of MCF7 cells data set of CMap. The MCF7 cells data set of CMap was subjected to clustering analysis with the ward method (Chiba & Takahashi, 1994). An arrow indicates the cluster where Hsp90 inhibitors belong. The numbers following the compound names indicate the ordinal numbers from the left.

Figure EV5. Decomposition of topoisomerase-inhibitor effect

A. Analysis of P17 factor using the method described in Fig 2D. Green or light salmon indicates topoisomerase inhibitors.

B. GO analysis of the P15 and P16 factors mainly constituting genes using the method described in Fig 2C. The obtained p-values were processed using the Benjamini-Hochberg method for multiple-testing corrections in each factor (α < 0.05). As for the P15 factor, GO terms associated with mitochondria are filled with yellow.

C. Analysis of the P15 factor using the method described in Fig 2D. Green indicates daunorubicin, doxorubicin, and mitoxantrone.

D. Analysis of the P16 factor using the method described in Fig 2D. Green indicates daunorubicin, doxorubicin, and mitoxantrone.

### Figure EV6. Identification of autophagy regulator

A. Analysis of the P2 factor using the method described in Fig 2D. Green or light salmon indicates PI3K/Akt/mTOR pathway inhibitors.

B. Clustering analysis of MCF7 cells data set of CMap. The MCF7 cells data set of CMap was subjected to clustering analysis with the ward method. The arrow indicates the cluster where the indicated compounds belong. The numbers following the compound names indicate the ordinal numbers from the left.

### Figure EV7. Relationship between contribution of factors and GO

A. A 2 × 2 contingency table for factors in CMap data set. “< 60%” and “> 60%” indicate the factors whose cumulative contribution is less and more than 60%, respectively. “GO (+)” or “GO (-)” represents the factors significantly annotated with or without GO. “P”, Fisher’s exact test probability.

B. A 2 × 2 contingency table for factors in BMDM data set. “< 60%” and “> 60%” indicate the factors whose cumulative contribution is less and more than 60%, respectively. “GO (+)” or “GO (-)” represents the factors significantly annotated with or without GO. “P”, Fisher’s exact test probability.

### Figure EV8. Identification of autophagy regulator

Western blotting analysis of HeLa cells treated with PP. HeLa cells were treated with PP at the indicated concentration for 6 h. Sirolimus is employed as a positive control for mTORC1 inhibition. The whole-cell lysate was analyzed by western blotting using anti-S6K and -P-S6K (Thr389) antibodies. A representative result of two independent experiments is shown.

## Abbreviations

AhR: aryl hydrocarbon receptor
BEN: benzamil
BC: benzethonium chloride
BMDM: bone marrow derived macrophages
BSA: bovine serum albumin
CMap: connectivity map
DMEM: Dulbecco’s Modified Eagle’s Medium
FA: factor analysis
FBS: fetal bovine serum
GO: gene ontology
IQ: 2-amino-3-methylimidazo[4,5-/]quinolone
LINCS: library of integrated network-based cellular signatures
LPS: lipopolysaccharide
MET: metiothepin
MEX: metixene
MOA: mode of action
mTORC: mammalian target of rapamycin complex
OLSA: orthogonal linear separation analysis
PCA: principal component analysis
PP: phenazopyridine
SEGR: significant enrichment of GO ratio
TCDD: 2,3,7,8-tetrachlorodibenzo-p-dioxin
WB: western blotting

